# Structure-guided bifunctional molecules hit a DEUBAD-lacking hRpn13 species upregulated in multiple myeloma

**DOI:** 10.1101/2021.07.16.452547

**Authors:** Xiuxiu Lu, Venkata R. Sabbasani, Vasty Osei-Amponsa, Christine N. Evans, Julianna C. King, Sergey G. Tarasov, Marzena Dyba, King C. Chan, Charles D. Schwieters, Sulbha Choudhari, Caroline Fromont, Yongmei Zhao, Bao Tran, Xiang Chen, Hiroshi Matsuo, Thorkell Andresson, Raj Chari, Rolf E. Swenson, Nadya I. Tarasova, Kylie J. Walters

## Abstract

Proteasome substrate receptor hRpn13 is a promising anti-cancer target. By integrated *in silico* and biophysical screening, we identified a chemical scaffold that binds hRpn13 with non-covalent interactions that mimic the proteasome and a weak electrophile for Michael addition. hRpn13 Pru domain binds proteasomes and ubiquitin whereas its DEUBAD domain binds deubiquitinating enzyme UCHL5. NMR revealed lead compound XL5 to interdigitate into a hydrophobic pocket created by lateral movement of a Pru β-hairpin with an exposed end for Proteolysis Targeting Chimeras (PROTACs). Implementing XL5-PROTACs as chemical probes identified a DEUBAD-lacking hRpn13 species (hRpn13^Pru^) present naturally with cell type-dependent abundance. XL5-PROTACs preferentially target hRpn13^Pru^, causing its ubiquitination. Gene-editing and rescue experiments established hRpn13 requirement for XL5-PROTAC-triggered apoptosis and increased p62 levels. These data establish hRpn13 as an anti-cancer target for multiple myeloma and introduce an hRpn13-targeting scaffold that can be optimized for preclinical trials against hRpn13^Pru^-producing cancer types.

## Introduction

The 26S proteasome is formed by a regulatory particle (RP) that binds and processes ubiquitinated substrates and a core particle (CP) that hydrolyzes proteins into peptides^1^. CP inhibitors are used to treat hematological cancers but resistance mechanisms motivate new strategies for proteasome inhibition^2^. Proteasome substrates are marked by covalently attached ubiquitin chains^3^ and the therapeutic potential of the ubiquitin-proteasome pathway in cancer treatment has exploded with new possibilities by invoking Proteolysis Targeting Chimeras (PROTACs), which link molecular targets to ubiquitination machinery^4^.

Rpn1, Rpn10, and Rpn13 in the RP bind ubiquitin or a shuttle factor carrying ubiquitinated substrates^5–13^ as well as ubiquitin-processing enzymes; namely, deubiquitinating enzymes UCHL5/Uch37^14–16^ and Usp14/Ubp6^17, 18^ for hRpn13 and hRpn1 respectively and E3 ligase E6AP/UBE3A for hRpn10^19^. The abundance of UCHL5^15, 20^ and E6AP^19^ in cells depends on their binding partners hRpn13 and hRpn10, respectively. The proteasome RP also has an essential deubiquitinating enzyme, Rpn11^21^, positioned near the substrate entrance that couples removal of ubiquitin chains with substrate translocation through the center of the proteasome ATPase ring by direct interaction with substrate-conjugated ubiquitin chains^22, 23^. Rpn11 interaction with ubiquitin chains at the proteasome does not require substrate^24^; thus, it likely plays an active role in positioning ubiquitinated substrates proximal to the nearby ATPase ring. Inhibitors against Rpn11 block cancer cell proliferation, induce the unfolded protein response, and/or trigger apoptosis^25, 26^.

CRISPR-based gene editing indicated hRpn13-binding compounds (RA190 and RA183)^27, 28^ to induce apoptosis in an hRpn13-dependent manner^29, 30^, albeit knockdown experiments suggest little dependency^31^, including for an hRpn13-binding peptoid^32, 33^. The C-terminal end of proteasome subunit hRpn2 extends across the hRpn13 N-terminal Pru (<u>P</u>leckstrin-like receptor for <u>u</u>biquitin) domain^34–36^ which also binds ubiquitin^6, 7^ dynamically, maintaining it in an extended conformation, with interactions at the ubiquitin linker region that cause preference for chains linked by K48^37, 38^. RA190 and RA183 react with hRpn13 C88 at the periphery of the hRpn2-binding region^27, 28^, but are generally reactive with exposed cysteines, impairing specificity^28, 31, 35^. Aided by a pipeline extending from *in silico* and biophysical integrated screening to high resolution structure determination, we generated bifunctional PROTAC-fused hRpn13-targeting compounds that require an intact hRpn13 Pru domain to induce apoptosis in multiple myeloma cells. We further used a lead compound (XL5-PROTAC) as a chemical probe of hRpn13 function and processing in cells.

## Results

### Structure-based screen finds an hRpn13-binding compound

We conducted *in silico* docking screens of commercial libraries containing 63 million compounds by using the hRpn13 Pru:hRpn2 structure^35, 36^ and hRpn2-binding site of hRpn13 as a binding pocket. Twenty-two potential lead compounds were selected for validation by biophysical assays (Supplementary Table 1). Binding to hRpn2 causes partially exposed W108^6, 39^ to be buried^35, 36^, allowing tryptophan quenching by differential scanning fluorimetry (DSF at λ_350_) as an indicator of binding. This approach was used to experimentally validate compound binding to the hRpn2-binding surface of hRpn13. In separate experiments for each of the twenty compounds, 20 μM compound was incubated with 1 μM hRpn13 Pru and fluorescence emission at 350 nm measured. Greatest tryptophan quenching was observed by XL5 addition (Supplementary Table 1) and incremental titration of XL5 into 1 μM hRpn13 Pru revealed concentration dependency (Fig. 1a). Ten candidate compounds, including XL5, were evaluated further by NMR; XL4, which demonstrated tryptophan quenching (Supplementary Table 1), was excluded by insolubility at the required concentration. The compounds were separately added at 10-fold molar excess to 20 μM ^15^N-labeled hRpn13 Pru and binding assessed at 25°C by 2D NMR for samples dissolved in NMR buffer (20 mM sodium phosphate, 50 mM NaCl, 2 mM DTT, 10% DMSO-*d*_6_ (deuterated DMSO), pH 6.5). XL5 and no other tested compound indicated binding to hRpn13 by 2D NMR. XL5 addition caused hRpn13 signals to shift from free state positions to an observable bound state whereas spectral changes were not induced by the other compounds tested (Supplementary Fig. 1a). Binding was also observed at 10°C with XL5 at 2-fold molar excess and hRpn13 at 0.25 mM (Fig. 1b); greater sample stability was observed at this lower temperature which was therefore used for the NMR experiments described below.

**Fig. 1.**
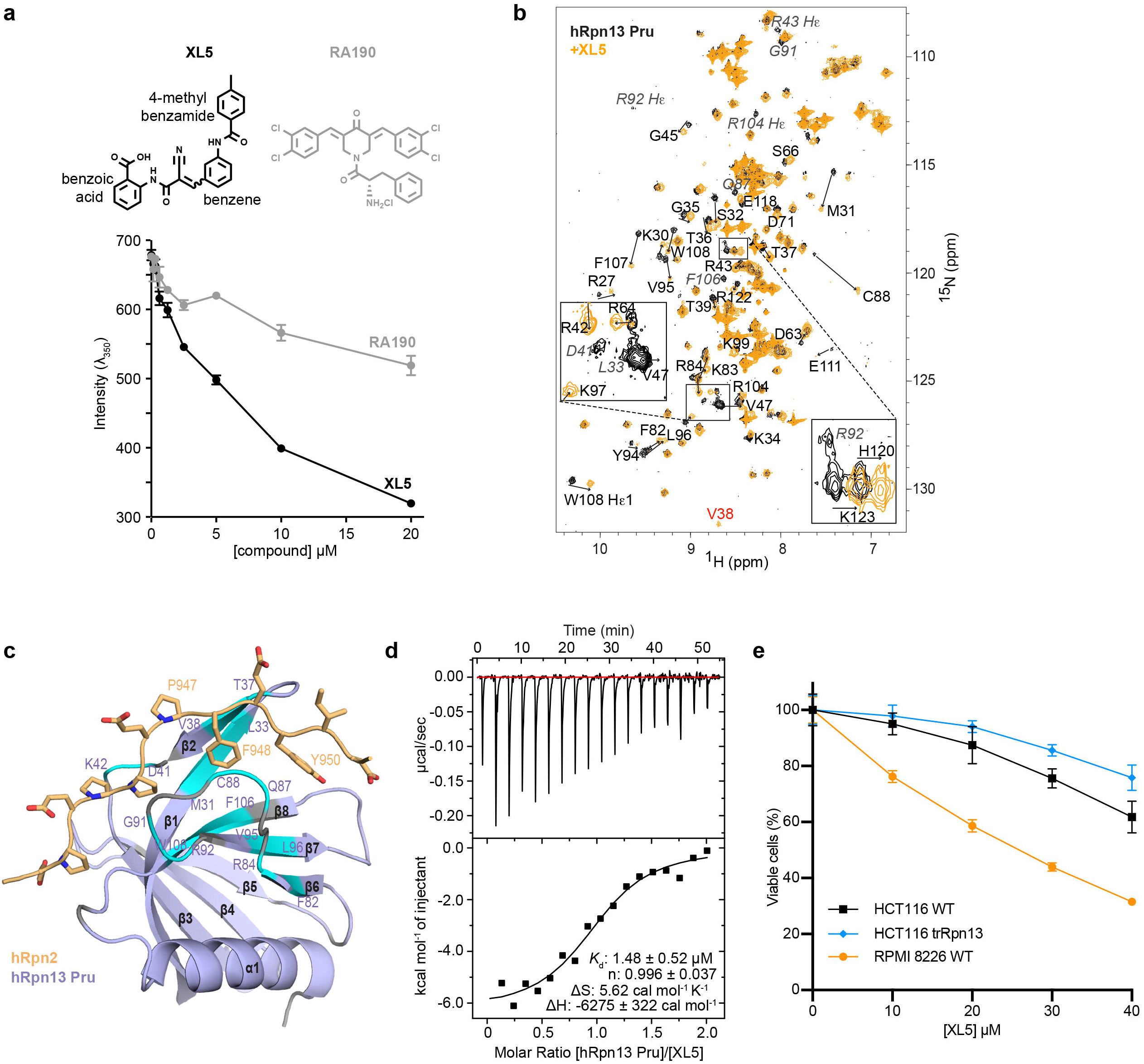
Structure-based screen yields an hRpn13-binding compound. **a**, Emission at 350 nm for 1 μM hRpn13 Pru with addition of XL5 (black) or RA190 (grey). The plots depict mean ± SD from three parallel recordings above which chemical structures are included. **b**, ^1^H, ^15^N HSQC spectra of 20 μM ^15^N-hRpn13 Pru (black) or 250 μM ^15^N-hRpn13 Pru with 2-fold molar excess XL5 (orange) in NMR buffer at 10°C, with an expansion for clarity. Arrows highlight the shifting of hRpn13 signals from their free state to their XL5-bound state. Residue signals that disappear (italicized grey) or V38 (red), which appears, following XL5 addition are labeled. **c**, hRpn13 amino acids significantly affected by XL5 addition in (**b**) are highlighted (light blue) on a secondary structure diagram of the hRpn13 Pru (purple):hRpn2 (940-953) (light orange) complex (PDB 6CO4). hRpn13 residues shifted by greater than one standard deviation above average or that appear (V38) or disappear following XL5 addition are highlighted. Prolines or residues not observed for free and XL5-bound hRpn13 are colored grey; hRpn2 side chain heavy atoms are displayed with nitrogen and oxygen colored blue and red respectively. **d**, ITC analysis of hRpn13 binding to XL5. Raw ITC data (top) from titration of 200 μM hRpn13 Pru into 20 μM XL5 with the binding isotherm and fitted thermodynamic values (bottom). **e**, HCT116 WT (black), HCT116 trRpn13 (blue) or RPMI 8226 WT (orange) cells were treated with the indicated concentration of XL5 in sextuplicate for 48 hours and cell metabolism measured by an MTT assay (mean ± SD). Viability is calculated as (l_570_)_sample_/(l_570_)_control_*100 (%).

Consistent with the tryptophan quenching detected by DSF (Fig. 1a), XL5 caused the epsilon and amide signals for W108 to shift (Fig. 1b). We quantified the shifting of the NMR signals following XL5 addition across the hRpn13 sequence to identify all significantly affected amino acids (Supplementary Fig. 1b). In some cases, signals appear or disappear, such as the V38 amide signal, which appears upon XL5 addition, or the amide signals for L33, D41, Q87, G91, R92, and F106 and epsilon signals for R43, R92, and R104, all of which disappear following XL5 addition (Fig. 1b). We mapped the hRpn13 amino acids most affected by XL5 onto a ribbon diagram of hRpn2-bound hRpn13 Pru (PDB 6CO4)^35^. The affected amino acids center around the region bound by hRpn2 F948 (Fig. 1c), which is required for hRpn2 binding to hRpn13^34^.

Isothermal titration calorimetry (ITC) was used to measure the binding affinity between hRpn13 and XL5. hRpn13 Pru was added incrementally to XL5 and the data fit to a 1-site binding mode (Fig. 1d). An overall binding affinity (*K*_d_) of 1.48 ± 0.52 μM was determined with favorable enthalpy and entropy. We attempted to measure the binding affinity of RA190 for the hRpn13 Pru by ITC but did not detect binding by this method, which relies on enthalpic changes (heat effects) (Supplementary Fig. 1c). Tryptophan fluorescence emission quenching was sensitive to RA190 addition to hRpn13 Pru, with a titration-dependent reduction in λ_350_ signal (Fig. 1a and Supplementary Table 1).

### XL5 treatment reduces viability of multiple myeloma cells

We tested whether XL5 restricts viability of RPMI 8226 multiple myeloma and HCT116 colon cancer cell lines by measuring metabolism with an MTT (3-(4,5-dimethylthiazol-2-yl)-2,5-diphenyltetrazolium bromide) assay. Experiments were also conducted in parallel with the HCT116 trRpn13 colon cancer cell line that expresses a truncated hRpn13 protein with a defective Pru and inability to bind the proteasome^30^. WT (wild-type) RPMI 8226 and WT or trRpn13 HCT116 cells seeded at 8,000 and 4,000 cells per well were treated with varying concentrations of XL5 extending to 40 μM and compared to cells incubated with equivalent amounts of DMSO vehicle control. Reduced metabolic activity was observed with XL5 treatment in a concentration-dependent manner for the two WT cell lines but higher concentration was required for HCT116 cells compared to RPMI 8226 cells (Fig. 1e). HCT116 cells were relatively insensitive to XL5, but potency was reduced in HCT116 trRpn13 cells (Fig. 1e).

### XL5 binds covalently to hRpn13 Pru

To define how hRpn13 interacts with XL5, we recorded unambiguous NOE interactions between hRpn13 and XL5 by acquiring a 3-dimensional ^1^H, ^13^C half-filtered NOESY experiment on a sample of ^13^C-labeled hRpn13 Pru mixed with 2-fold molar excess unlabeled XL5 (Fig. 2a-b). Chemical shift values were assigned to hRpn13 and XL5 (Fig. 2a-b, Supplementary Fig. 2 and Supplementary Table 2) as described in Methods. Protons indicating saturation of the XL5 alkene group (H13 and H19 in Fig. 2a, left panel) were present in the spectrum forming NOE interactions with methyl groups of hRpn13 V85 and V93 (Fig. 2a, right panel and Supplementary Table 3). These interactions suggested that XL5 interacts with hRpn13 by Michael addition at C88, similar to RA190 (Fig. 1a) and consistent with the shifting of hRpn13 C88 observed in 2D NMR spectra following XL5 addition (Fig. 1b and Supplementary Fig. 1b). To test for a covalent interaction directly, an hRpn13 Pru sample was incubated with 10-fold molar excess XL5 or DMSO (vehicle control) and subjected to liquid chromatography-mass spectrometry (LC-MS). A product was detected of appropriate molecular weight for covalent addition of XL5 to hRpn13 Pru (Fig. 2c) that was absent from the control experiment albeit with low abundance (Supplementary Fig. 3a). This low abundance may be caused by the harsh experimental conditions of LC-MS and reversibility of the carbon-sulfur bond.

**Fig. 2.**
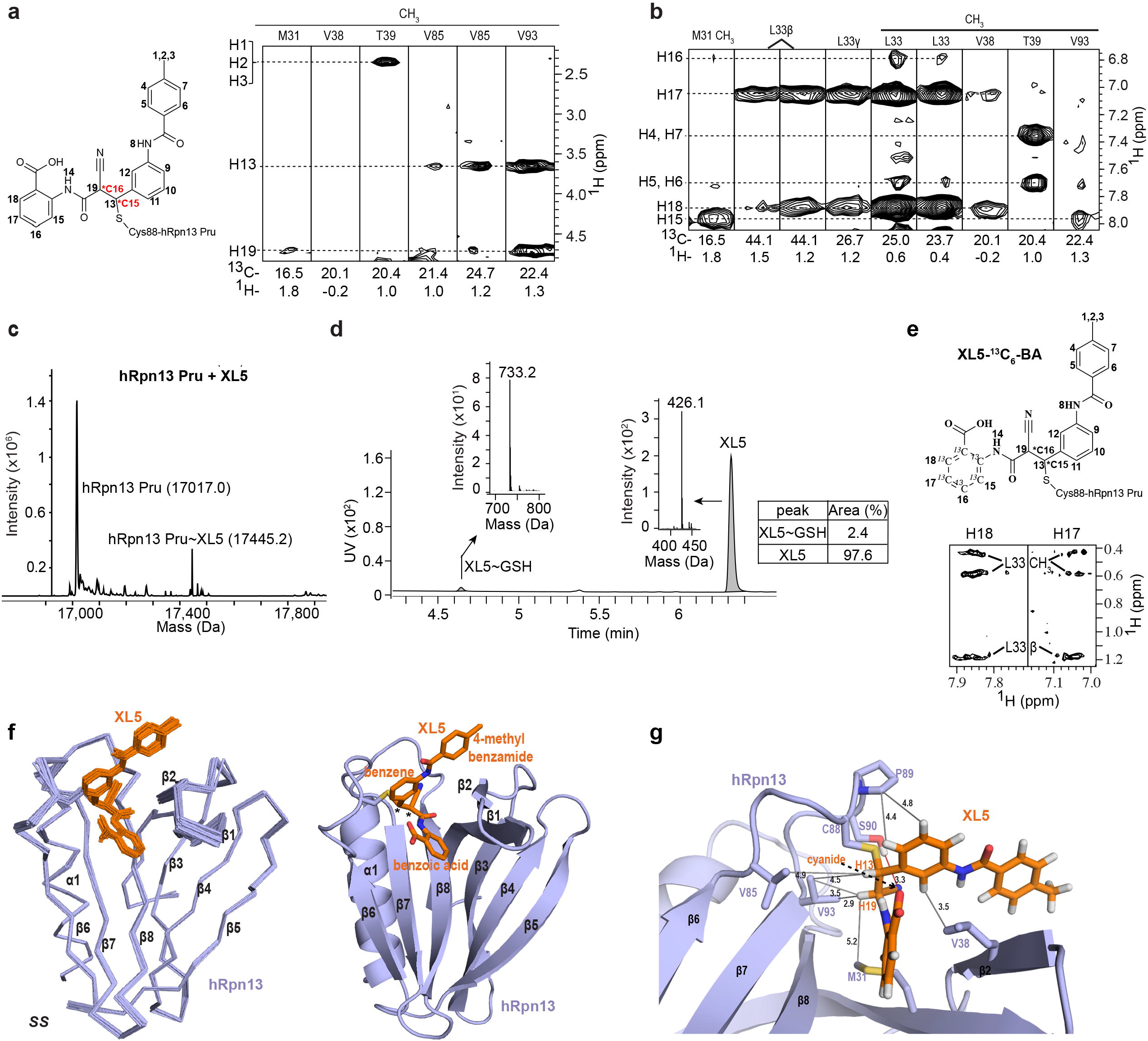
Structure of XL5-ligated hRpn13 Pru. **a-b**, Chemical structure of XL5 (left panel) including the ligated sulfur atom from hRpn13 C88. Hydrogen atoms are labeled with numbers used in the text and figures. Chiral center C15 or C16 is labeled and indicted with a star in red. Selected regions from a ^1^H, ^13^C half-filtered NOESY (100 ms) experiment (a, right panel and b) acquired on a sample containing 0.25 mM ^13^C-labeled hRpn13 Pru and 2-fold molar excess unlabeled XL5 dissolved in NMR buffer. **c**, LC-MS analysis of 2 μM purified hRpn13 Pru (MW: 17017.3 g/mol) incubated with 20 μM XL5 for 2 hours at 4°C. The resulting compound adduct and unmodified hRpn13 Pru are labeled along with the detected molecular weight (Da). **d**, LC-MS analysis of 40 μM XL5 incubated with 2 mM reduced L-glutathione (GSH, MW: 307.3 g/mol) for 2 hours at 4°C. Detected GSH adducts are indicated and a table is included that lists relative abundance. **e**, Chemical structure of XL5-^13^C_6_-BA (upper panel) illustrating ^13^C-labeling. Selected regions from a ^1^H, ^13^C half-filtered NOESY (100 ms) experiment (lower panel) acquired on a sample with 0.4 mM unlabeled hRpn13 Pru and equimolar of XL5-^13^C_6_-BA dissolved in NMR buffer containing 70% ^2^H_2_O. **f,** Structural ensemble (left panel) or ribbon diagram (right panel) of hRpn13 (purple) ligated to XL5 (orange) with C15 and C16 in the SS stereoconfiguration. hRpn13 secondary structural elements and XL5 chemical groups are labeled with the two chiral centers indicated by an asterisk (*). The C88 sulfur and XL5 nitrogen and oxygen atoms are colored yellow, blue and red respectively in the ribbon diagram. **g**, Enlarged view highlighting interactions between hRpn13 M31, V85 and V93 with XL5 H13 and H19 as well as hRpn13 V38 and P89 with the XL5 central benzene. A weak hydrogen bond is formed between the hRpn13 S90 hydroxy group and XL5 cyanide group (red line). Key interactions are highlighted (grey lines) including distances (Å) for XL5 hydrogen or cyanide nitrogen atoms with hRpn13 carbon atoms (colored as in **f** right panel).

We tested whether XL5 exhibits specificity for hRpn13 by incubating 40 μM XL5 at 4°C for two hours with 2 mM reduced L-glutathione, a non-protein thiol containing representative of non-specific interactions with exposed cysteines. XL5-ligated glutathione was detected at only 2% abundance (Fig. 2d). Under identical conditions, 40 μM RA190 reacted with 2 mM reduced L-glutathione to yield products with one or two molecules ligated at 14% or 30% abundance, respectively (Supplementary Fig. 3b). We also tested XL5 reactivity by incubating it at 0.2 µM with mouse serum (BioIVT) and monitoring stability by LC-MS over a 24-hour period to find only 6% reduction (Supplementary Fig. 3c).

### Structure of XL5-ligated hRpn13

Model structures predicted from the *in silico* screen conflicted with our experimental data, as these indicated XL5 to bind non-covalently to hRpn13 at a location somewhat different from that suggested by the NMR data (Supplementary Fig. 4). We therefore used NMR, including the ^1^H, ^13^C half-filtered NOESY experiment described above (Fig. 2a-b), to solve the structure of XL5-ligated hRpn13. NOEs involving XL5 H15-H18 were detected to hRpn13 methyl groups of M31, L33, V38 and V93 (Fig. 2b). These interactions were validated by selective ^13^C-labeling of the XL5 benzoic acid ring (Fig. 2e, XL5-^13^C_6_-BA). Mixing this isotopically labeled XL5 compound with equimolar unlabeled hRpn13 Pru revealed in a 3-dimensional ^1^H, ^13^C half-filtered NOESY experiment NOEs between hRpn13 L33 and XL5 H17 and H18 (Fig. 2e, bottom panel); the weaker interactions involving hRpn13 V38 as well as XL5 H15 and H16 (Fig. 2b) were not observable in this less sensitive experiment. Signals from H4 and H5 of the XL5 4-methyl benzamide group are indistinguishable compared to H7 and H6 respectively (Fig. 2a, left panel), but interactions were recorded between H4/H7 and H5/H6 of XL5 and hRpn13 T39 (Fig. 2b), which also exhibited NOE interactions with the XL5 methyl group (Fig. 2a, right panel). In total, we recorded 23 NOE interactions between hRpn13 and XL5 (Fig. 2a-b, Table 1 and Supplementary Table 3).

**Table 1.**
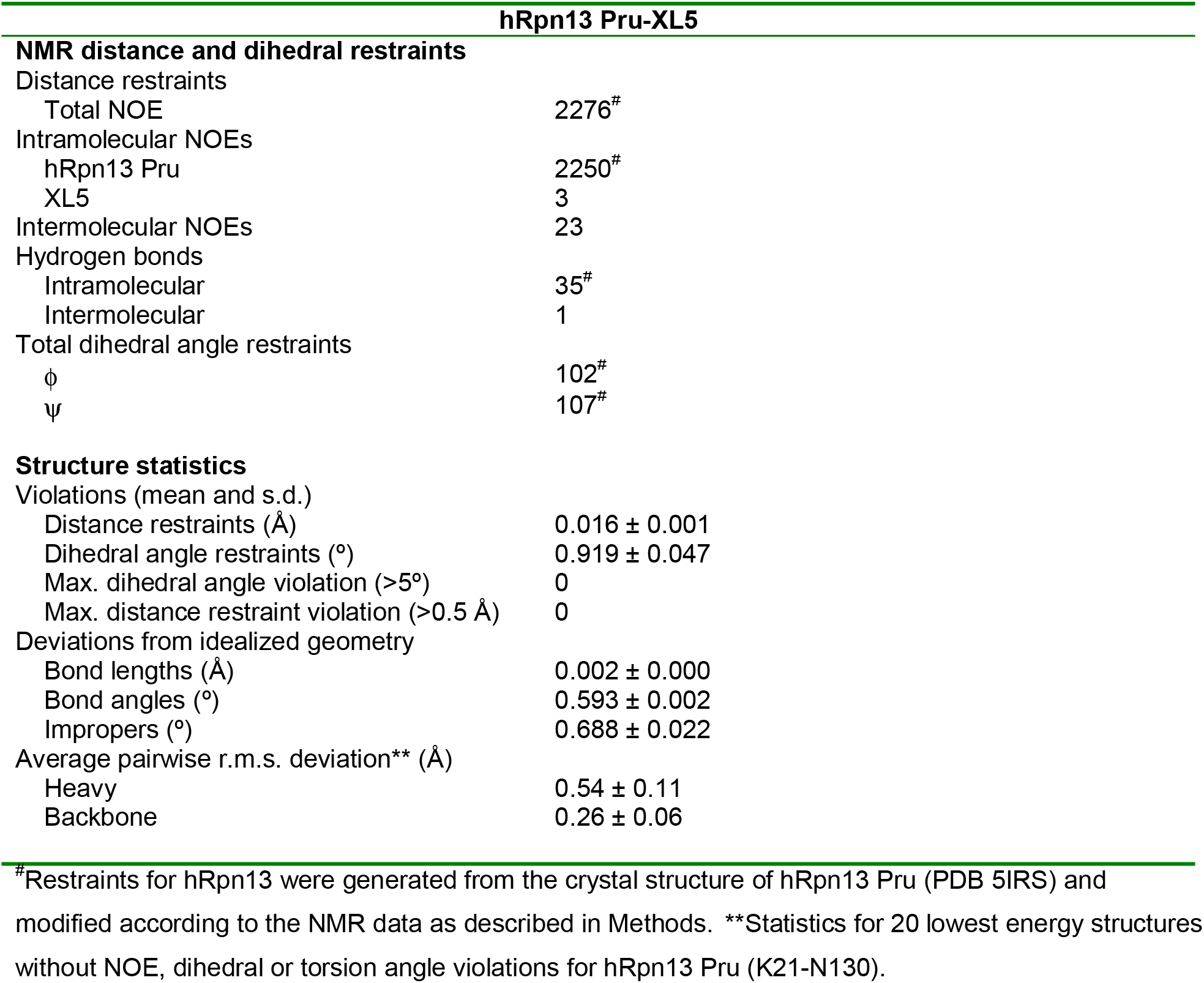
NMR and refinement statistics for XL5-ligated hRpn13 Pru.

| hRpn13 Pru-XL5 |  |
| --- | --- |
| **NMR distance and dihedral restraints** |  |
| Distance restraints |  |
| Total NOE | 2276<sup>#</sup> |
| Intramolecular NOEs |  |
| hRpn13 Pru | 2250<sup>#</sup> |
| XL5 | 3 |
| Intermolecular NOEs | 23 |
| Hydrogen bonds |  |
| Intramolecular | 35<sup>#</sup> |
| Intermolecular | 1 |
| Total dihedral angle restraints |  |
| $\phi$ | 102<sup>#</sup> |
| $\psi$ | 107<sup>#</sup> |
| **Structure statistics** |  |
| Violations (mean and s.d.) |  |
| Distance restraints (Å) | 0.016 ± 0.001 |
| Dihedral angle restraints (°) | 0.919 ± 0.047 |
| Max. dihedral angle violation (>5°) | 0 |
| Max. distance restraint violation (>0.5 Å) | 0 |
| Deviations from idealized geometry |  |
| Bond lengths (Å) | 0.002 ± 0.000 |
| Bond angles (°) | 0.593 ± 0.002 |
| Impropers (°) | 0.688 ± 0.022 |
| Average pairwise r.m.s. deviation** (Å) |  |
| Heavy | 0.54 ± 0.11 |
| Backbone | 0.26 ± 0.06 |

<sup>#</sup>Restraints for hRpn13 were generated from the crystal structure of hRpn13 Pru (PDB 5IRS) and modified according to the NMR data as described in Methods. \*\*Statistics for 20 lowest energy structures without NOE, dihedral or torsion angle violations for hRpn13 Pru (K21-N130).

When ligated to hRpn13 C88, XL5 C15 and C16 (Fig. 2a, left panel) can in principle adopt either R or S stereochemistry and we therefore initially calculated structures for XL5-ligated hRpn13 with all possible stereochemistry, including SS, RR, SR and RS for C15 and C16 respectively. As discussed in Methods, only SS stereochemistry fit the NOESY data (Supplementary Fig. 5) although it remains possible a population exists that includes R stereochemistry at either of these two sites but with too low of an abundance for NOE detection. The calculated structures for SS stereochemistry converged with a heavy atom root-mean-square-deviation (r.m.s.d.) of 0.54 Å (Fig. 2f, left panel and Table 1). A key feature of XL5 interaction with hRpn13 is the sulfide bond formed to the C88 thiol group (Fig. 2f, right panel and 2g, yellow) facilitated by nearby interactions from XL5 H13 and H19 to hRpn13 M31, V85, and V93 methyl groups (Fig. 2g); these interactions are dictated by the NOESY data (Fig. 2a, right panel).

### Structure validation by chemical probing reveals a site for PROTAC addition

The overall structure of hRpn13 ligated to XL5 is similar to the unligated (PDB 5IRS)^9^ (Fig. 3a) and hRpn2-bound (PDB 6CO4)^35^ (Fig. 3b-c) structures, as expected from the NOEs detected within the structural core in a 3-dimensional ^13^C-dispersed NOESY experiment acquired on ^13^C-labeled hRpn13 Pru mixed with 1.2-fold molar excess unlabeled XL5 (Supplementary Fig. 6). To accommodate XL5, however, the hRpn13 β1-β2 hairpin is shifted away from β8 (Fig. 3a-c), allowing intercalation of the benzoic acid group within a hydrophobic pocket formed by β1 L33, β2 V38, and β8 F106 (Fig. 3b). In the XL5-ligated structure, hRpn13 W108 Hβ and Cγ are close to XL5 H19 and the cyanide group (Fig. 3a and Supplementary Table 4). These interactions coupled with the change in chemical environment of W108 due to the reconfiguration of local structure (Fig. 3a) provides an explanation for its observed Hε1 and amide resonance shifting (Fig. 1b and Supplementary Fig. 1a-b) and reduction of intrinsic emission at λ_350_ (Fig. 1a and Supplementary Table 1).

**Fig. 3.**
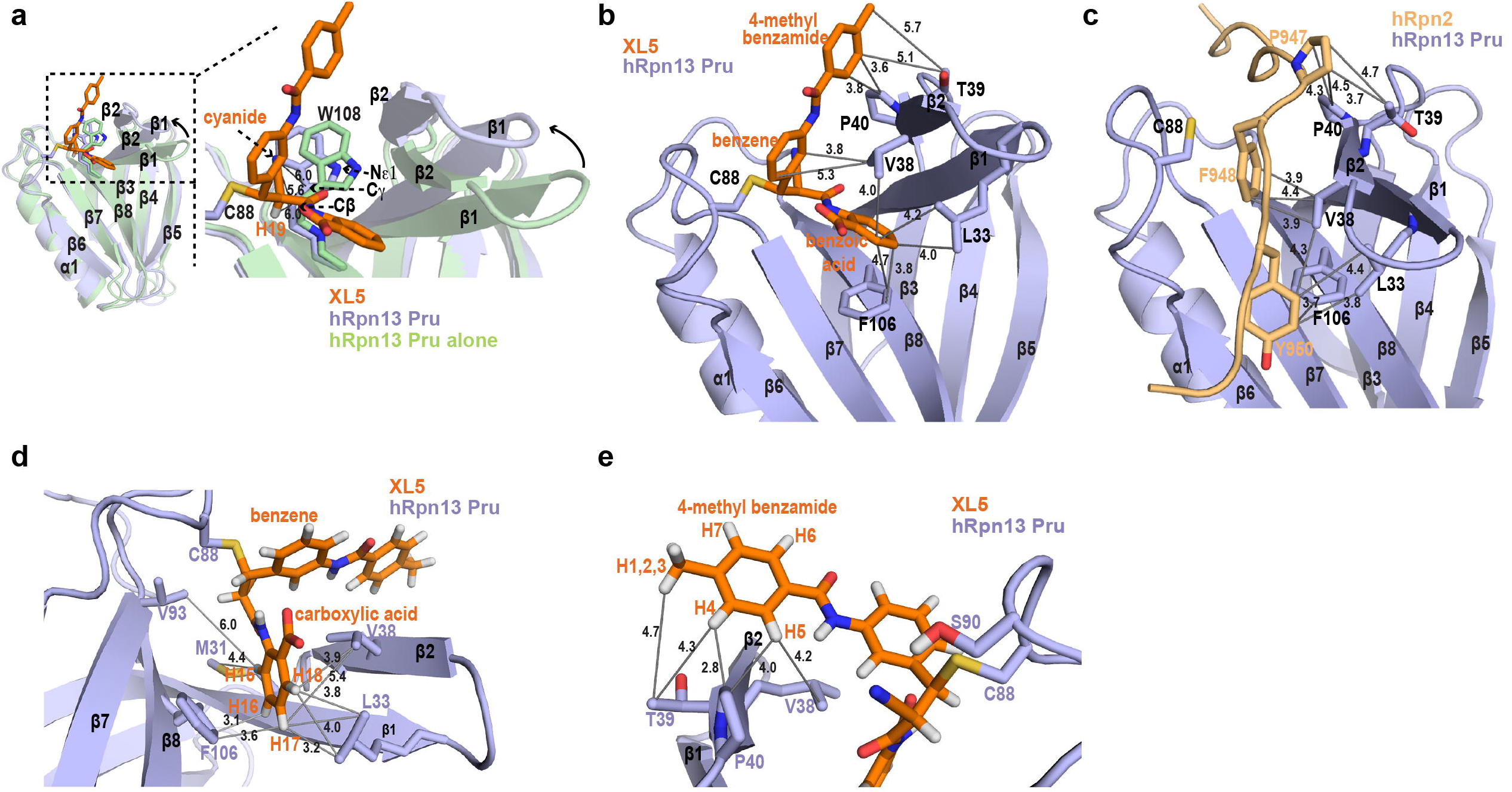
hRpn13 β-hairpin shifts to bury XL5 benzoic acid. **a-e,** Ribbon diagram structures of hRpn13 Pru ligated to XL5 (colored as in Fig. 2g) to highlight key interactions, which are indicated by grey lines with distances (Å) included. **a**, Comparison of XL5-ligated and free hRpn13 Pru (PDB 5IRS, green) structures with an expansion (dashed rectangles) in the right panel and hRpn13 W108 included. **b-c**, Structural comparison of XL5-ligated hRpn13 (colored as Fig. 2g) and hRpn2-bound hRpn13 (PDB: 6CO4) with hRpn2 colored as in Fig. 1c. **d**, Expanded view highlighting hRpn13 M31, L33, V38, and V93 interaction with the XL5 benzoic acid group. **e**, Expanded view of XL5 4-methyl benzamide interaction with hRpn13 V38, T39 and P40.

XL5 binds to hRpn13 Pru with a similar affinity as hRpn2 (944-953)^35^ and forms analogous interactions. The central aromatic ring is positioned close to where hRpn2 F948 binds and similarly interacts with V38 while the XL5 4-methyl benzamide binds hRpn13 T39 and P40 similarly compared to hRpn2 P947 (Fig. 3b-c). The shorter distance between the central benzene and benzoic acid groups of XL5 relative to hRpn2 F948 and Y950 (which are separated by E949) alters interactions with hRpn13 L33, V38 and F106 causing this end of XL5 to be buried (Fig. 3b-c). Consistent with this burying of the benzoic acid aromatic ring (Fig. 3d), inclusion of additional chemical groups to the XL5 scaffold caused reduced affinity (Supplementary Table 5). A bulky ortho-trifluoromethyl group (XL30 in Supplementary Table 5) caused ∼8-fold reduced affinity; this group would form steric clashes with the L33 methyl groups if bound in the same configuration as XL5. Reduced affinity was similarly caused by addition of methoxy (XL28) or methylamino (XL29) groups at the meta position (Supplementary Table 5 and Supplementary Fig. 7).

As XL5 H13 and H19 are directed towards the β6-β7 loop, the cyanide group is positioned to form a weak hydrogen bond to the hRpn13 S90 hydroxy group, placing the central benzene ring proximal to V38 and P89 (Fig. 2g). Addition of a trifluoromethyl group (Supplementary Table 5, XL32) or methylamino group (Supplementary Table 5, XL33) at either ortho position of the XL5 central benzene ring reduced binding affinity to hRpn13 (Supplementary Fig. 7) and the structure suggests that this reduction is due to steric clashes with C88 or V38, respectively. NMR signals of the central XL5 benzene ring are absent, which is consistent with the anion-π interaction formed between the XL5 π carboxylic acid group and central benzene (Fig. 3d); a similar broadening mechanism is reported for an anion (fluoride)-π (thiophene) interaction system. Replacement of this ortho carboxyl group with sulfonamide (XL31) strongly reduced affinity for hRpn13, potentially due to weakening of the XL5 anion-π interaction (Fig. 3d and Supplementary π Fig. 7). This part of the structure is well-defined (Fig. 3d) by NOE interactions observed to each end of XL5 as well as to H13 and H19 (Fig. 2a-b).

XL5 4-methyl benzamide interacts with the C-terminal end of hRpn13 β2 through hydrophobic interactions (Fig. 3d-e), which are indicated in the NOESY data (Fig, 2a-b). Modification of the 4-methyl benzamide ring to less hydrophobic 6-hydroxy-5-methyl-pyridine (XL27) reduced affinity compared to XL5 (Fig. 3e, Supplementary Table 5 and Supplementary Fig. 7), demonstrating the importance of these interactions. The XL5 4-methyl benzamide aromatic ring interacts with the β2 V38 methyl group that is close to the central benzene and P40. The methyl group interacts favorably with that of hRpn13 T39 (Fig. 3e) and its removal in XL23, coupled with inclusion of an ortho-chlorine, reduces affinity by >2-fold and substitution with trifluoromethyl (XL26) or carboxymethyl amino (XL25) groups similarly reduced affinity for hRpn13 Pru (Supplementary Table 5 and Supplementary Fig. 7). Substitution of the methyl group however with a methylamino group (XL24) had little effect (Supplementary Table 5 and Supplementary Fig. 7), which as described below, led us to use this site for PROTAC addition.

### Engineered cell lines establish hRpn13 requirement for XL5-PROTAC-induced apoptosis

Based on the structure and chemical probing described above, we extended XL5 at the methyl group position to include either of three established PROTACs, namely Von-Hippel Lindau (VHL, with two different linkers to XL5 and in one case, a VHL variation^41^), cereblon (CRBN) or inhibitor of apoptosis (IAP) (Fig. 4a). An MTT assay demonstrated greater cellular sensitivity when XL5 was fused to a PROTAC (Fig. 4b) with the hook effect^4^ observed for cells treated with XL5-VHL-2. Control reagents VHL ligand and thalidomide (for cereblon) did not affect metabolic activity even at 40 μM treatment; however, RPMI 8226 cells were sensitive to IAP ligand (Fig. 4a-b and Supplementary Fig. 8), which is reported to induce apoptosis^42^.

**Fig. 4.**
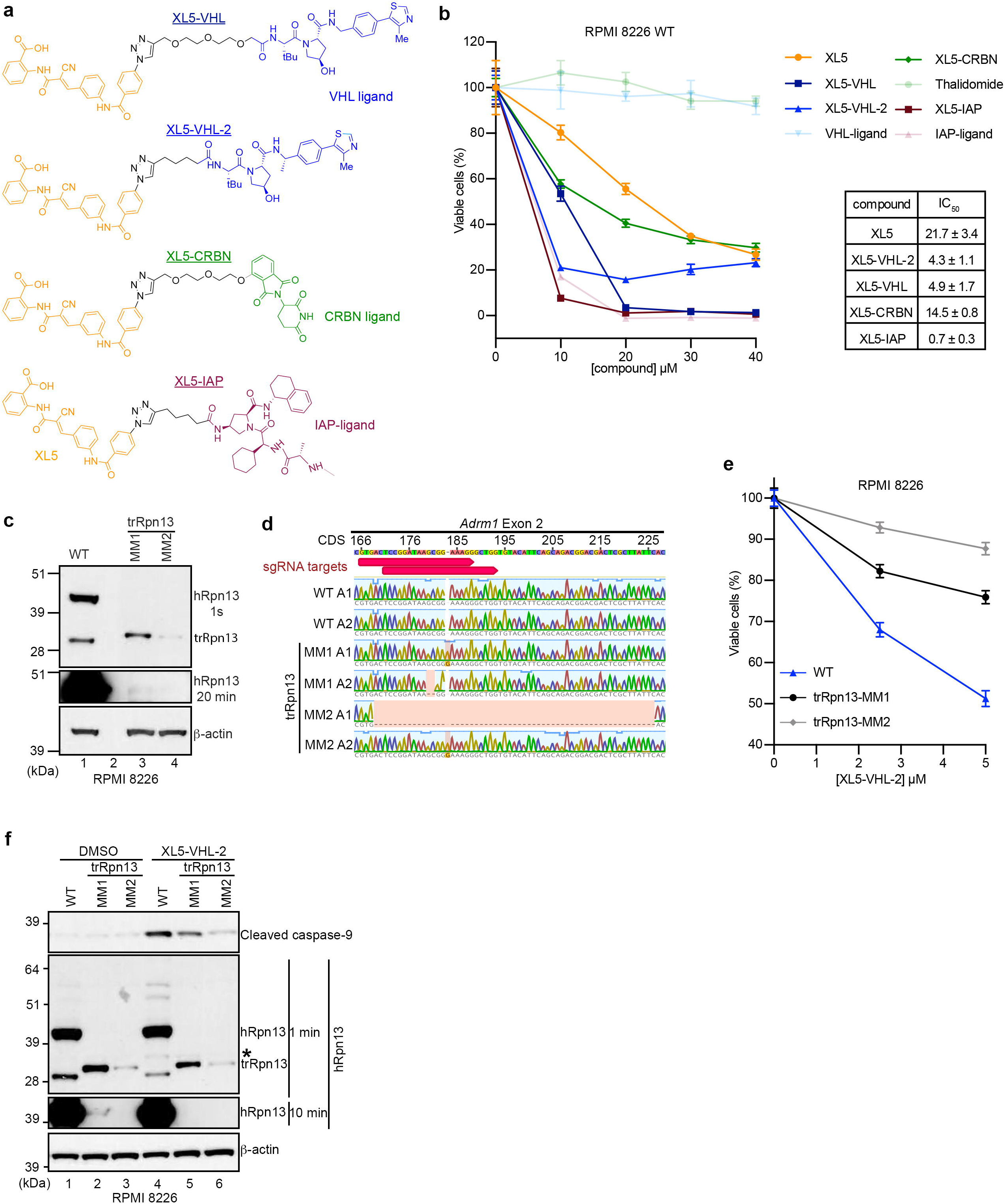
Engineered cell lines establish hRpn13 requirement for XL5-PROTAC-induced apoptosis. **a**, Chemical structures of XL5 (orange)-PROTACs (VHL, blue; CRBN, green, IAP, burgundy). **b**, RPMI 8226 cells were treated with the indicated concentration of XL5 (orange), XL5-VHL (navy), XL5-VHL-2 (blue), XL5-CRBN (green), XL5-IAP (burgundy), VHL-ligand (light blue), thalidomide (light green) or IAP-ligand (pink) in sextuplicate for 48 hours and cell metabolism measured by an MTT assay (mean ± SD). Viability is plotted as (l_570_)_sample_/(l_570_)_control_*100 (%). IC_50_ values are listed for XL5 and XL5-PROTAC. **c**, Immunoblot of whole cell extract from RPMI 8226 WT, trRpn13-MM1, or trRpn13-MM2 cells probing hRpn13 (1 second and 20 minutes exposure) or β-actin. **d**, Sanger sequencing analyses of hRpn13 cDNA from RPMI 8226 WT, trRpn13-MM1, or trRpn13-MM2 cells denoting the location of the two sgRNAs (red arrow) on hRpn13-encoding gene *Adrm1* Exon2 with cDNA sequence (CDS) labeled. Allele is abbreviated as “A”. **e**, RPMI 8226 WT (blue), trRpn13-MM1 (black) or trRpn13-MM2 (grey) cells were treated with the indicated concentrations of XL5-VHL-2 in sextuplicate for 48 hours and cell metabolism measured by an MTT assay (mean ± SD). Viability is calculated as (l_570_)_sample_/(l_570_)_control_*100 (%). **f,** Immunoblots of whole cell lysate from RPMI 8226 WT, trRpn13-MM1, or trRpn13-MM2 cells treated for 24 hours with 40 μM XL5-VHL-2 with comparison to DMSO (vehicle control) immunoprobing for cleaved caspase-9 (top panel), hRpn13 (two middle panels with 1 min or 10 min exposure), or β-actin (as a loading control, bottom panel). A black asterisk indicates cleaved caspase-9 in the 1-minute immunoblot for hRpn13, as hRpn13 was probed following cleaved caspase-9 and without stripping the membrane.

We next sought to test whether hRpn13 is the bone fide target of XL5-VHL-2. The comparison of HCT116 WT and trRpn13 cell lines is not a good measure of hRpn13 requirement as cell viability is not substantially impacted by XL5 treatment in the WT cell line (Fig. 1e). We therefore generated RPMI 8226 cells with the hRpn13 Pru disrupted by CRISPR/Cas9 gene editing with Exon 2 targeting of hRpn13-expressing *ADRM1* as done previously in HCT116 cells^30^. Putative edited cell lines were identified by targeted Illumina sequencing. Of 11 lines identified, two viable clones (trRpn13-MM1 and trRpn13-MM2) were validated and utilized for experiments. Immunoblotting the whole cell lysate with anti-hRpn13 antibodies revealed loss of full length hRpn13 with concomitant and differential expression of a truncated hRpn13 species at the molecular weight observed in HCT116 trRpn13 cells^30^ (Fig. 4c). Genomic sequencing indicated these two cell lines to each contain two alleles with a 50/50 split whereby one allele of Exon 2 had a one-nucleotide insertion and the other allele of trRpn13-MM1 and trRpn13-MM2 had a two and 58 nucleotide deletion respectively (Fig. 4d).

To test whether hRpn13 is required for XL5-VHL-2 cellular toxicity, we compared the effect of XL5-VHL-2 treatment for RPMI 8226 WT cells versus the two trRpn13-MM cell lines. Cellular metabolic activity was measured with an MTT assay, as done in Fig. 1e. The cell lines were seeded separately at 8,000 cells per well and treated with 2.5 or 5.0 μM concentration of XL5-VHL-2 or equivalent amounts of DMSO vehicle control. The potency of XL5-VHL-2 was reduced in both trRpn13-MM cell lines compared to WT RPMI 8226 cells (Fig. 4e). Surprisingly, trRpn13-MM1 was more sensitive to XL5-VHL-2 than trRpn13-MM2. The activity of XL5-VHL-2 was investigated further in these cell lines by directly probing for apoptosis with cleaved caspase-9 as an indicator. Each of the three RPMI 8226 cell lines (WT, trRpn13-MM1, trRpn13-MM2) were treated with 40 μM XL5-VHL-2 or DMSO (vehicle control) and immunoprobed for cleaved caspase-9. RPMI 8226 WT cells indicated the expected sensitivity to XL5-VHL-2 treatment (Fig. 4f, lane 4 versus lane 1). The two trRpn13-MM cell lines demonstrated reduced levels of cleaved caspase-9 compared to WT RPMI 8226 cells (Fig. 4f, lane 4, 5 and 6); however, as was observed for the MTT assay (Fig. 4e), the loss of XL5-VHL-2 potency was greater for trRpn13-MM2 (Fig. 4f). A longer exposure (ten minutes) of the membrane probed with anti-hRpn13 antibodies revealed low levels of full-length hRpn13 in trRpn13-MM1 but not trRpn13-MM2 (Fig. 4f, lane 2 and 3). We also observed low levels of hRpn13 in RPMI 8226 trRpn13-MM1 cells without loading samples from WT and trRpn13-MM1 cells next to each other (Fig. 4c, 20 min exposure for hRpn13, lane 1 versus 3) excluding the possibility of spillover occurrence (Fig. 4f, lane 1 versus 2). We next tested whether mRNA corresponding to the full length hRpn13 could be observed by PacBio sequencing on samples extracted from RPMI 8226 WT, trRpn13-MM1 and trRpn13-MM2 cells. Consistent with the immunoblotting (Fig. 4c and 4f), mRNA corresponding to full length hRpn13 was detected in RPMI 8226 WT and trRpn13-MM1 cells, but not trRpn13-MM2 cells (Supplementary Data 1). The abundance of full length hRpn13-encoding mRNA in trRpn13-MM1 was significantly reduced compared to WT (Supplementary Data 1, “FL” and “ORF_length” columns), consistent with the protein levels (Fig. 4c and 4f). The abundance of trRpn13 mRNA in trRpn13-MM2 cells was lower compared to trRpn13-MM1 cells (Supplementary Data 1, “FL” and “ORF_length” columns, 299 amino acids), corresponding to the lower protein levels of trRpn13 in trRpn13-MM2 cells (Fig. 4c and 4f). Although we do not know how trRpn13-MM1 cells transcribe full length hRpn13 mRNA, XL5-VHL-2-treatment led to clearance of hRpn13 full length protein from trRpn13-MM1 cells (Fig. 4f, lane 2 versus 5). The lower hRpn13 levels in trRpn13-MM1 appears to make it more sensitive to XL5-VHL-2-treament (Fig. 4f, lane 2 versus 5) as WT cells displayed similar hRpn13 levels, although upper molecular weight bands appeared between 51 and 64 kDa following XL5-VHL-2 treatment (Fig. 4f, lane 1 versus 4).

We previously found UCHL5 to be targeted by RA190^35^ and multiple studies have demonstrated the cellular abundance of UCHL5 to be dependent on hRpn13^15, 20, 30, 35^. To test whether UCHL5 is targeted by XL5-VHL-2, we immunoprobed the membrane from Fig. 4f for UCHL5. As observed in other cell lines^15, 20, 30, 35^, UCHL5 protein levels (Supplementary Fig. 9) correlated with hRpn13 protein levels (Fig. 4f) indicating greater reduction in trRpn13-MM2 than trRpn13-MM1; a caveat to this observation however is the presence of a likely non-specific band above UCHL5 that is unmodulated in WT compared to trRpn13-MM lines. Nonetheless, neither the UCHL5 band nor the band above it was changed by treatment with XL5-VHL-2. Moreover, no higher molecular weight UCHL5 species suggestive of ubiquitination were detected in XL5-VHL-2-treated RPMI 8226 WT cells compared to DMSO (control) (Supplementary Fig. 9, lane 1 versus 4).

Altogether, these experiments led to two conclusions; 1) the level of hRpn13 directly correlates with induction of apoptosis by XL5-VHL-2 and 2) substantial reduction in hRpn13 level retains of its role in induced apoptosis by XL5-VHL-2. This data thus highlights limitations of earlier studies that relied on incomplete knockdown by siRNA^31, 32^.

### Chemical probing reveals a DEUBAD-lacking hRpn13 species present with cell line dependency

To test further whether the XL5-PROTACs cause ubiquitination and/or degradation of hRpn13, lysates from WT RPMI 8226 cells treated with 40 μM XL5 or XL5-PROTAC were compared to DMSO control by immunoprobing for hRpn13 and β-actin loading control (Fig. 5a). The level of hRpn13 was similar in all treated RPMI 8226 cells (Fig. 5a, 1 second exposure); however, following longer exposure (3 minutes) of the membrane, an increase in higher molecular weight hRpn13 species characteristic of ubiquitination was observed for cells treated with XL5-VHL, XL5-VHL-2 or XL5-IAP with the two aforementioned bands between 51 and 64 kDa (Fig. 5a and 4f). In addition, a lower molecular weight band was observed that was correspondingly reduced in abundance following treatment with XL5-VHL, XL5-VHL-2, and XL5-IAP (Fig. 5a and 4f). RPMI 8226 cells are reported to be resistant to cereblon targeting^43–45^ and no effect was observed for hRpn13 or its lower molecular weight species following XL5-CRBN treatment (Fig. 5a). The antibody used to immunoprobe for hRpn13 recognizes an epitope that spans amino acids 100 – 200 (Abcam, personal communication), which includes a portion of the hRpn13 Pru and following interdomain linker region (Fig. 5b). To investigate whether the observed lower molecular weight hRpn13 species has an intact Pru, we tested whether it is pulled out of RPMI 8226 WT cells by GST-hRpn2 (940-953) encompassing the hRpn13-binding site at the proteasome^35^. Both hRpn13 full length protein and the smaller species bound to GST-hRpn2-bound glutathione Sepharose 4B resin and not to resin treated equivalently with GST control (Fig. 5c). We henceforth refer to this hRpn13 species as hRpn13^Pru^, as it contains an intact Pru domain. We next tested whether hRpn13^Pru^ is present at proteasomes of WT RPMI 8226 cells. Proteasomes were immunoprecipitated from whole cell lysates with anti-hRpt3 (a proteasome ATPase subunit) or rabbit IgG (as a control) antibodies and immunoprobed for hRpn13 as well as proteasome subunits hRpn2 and hRpt3, as controls. Both full length hRpn13 and hRpn13^Pru^ immunoprecipitated with hRpt3 (Fig. 5d), consistent with the pulldown experiment (Fig. 5c).

**Fig. 5.**
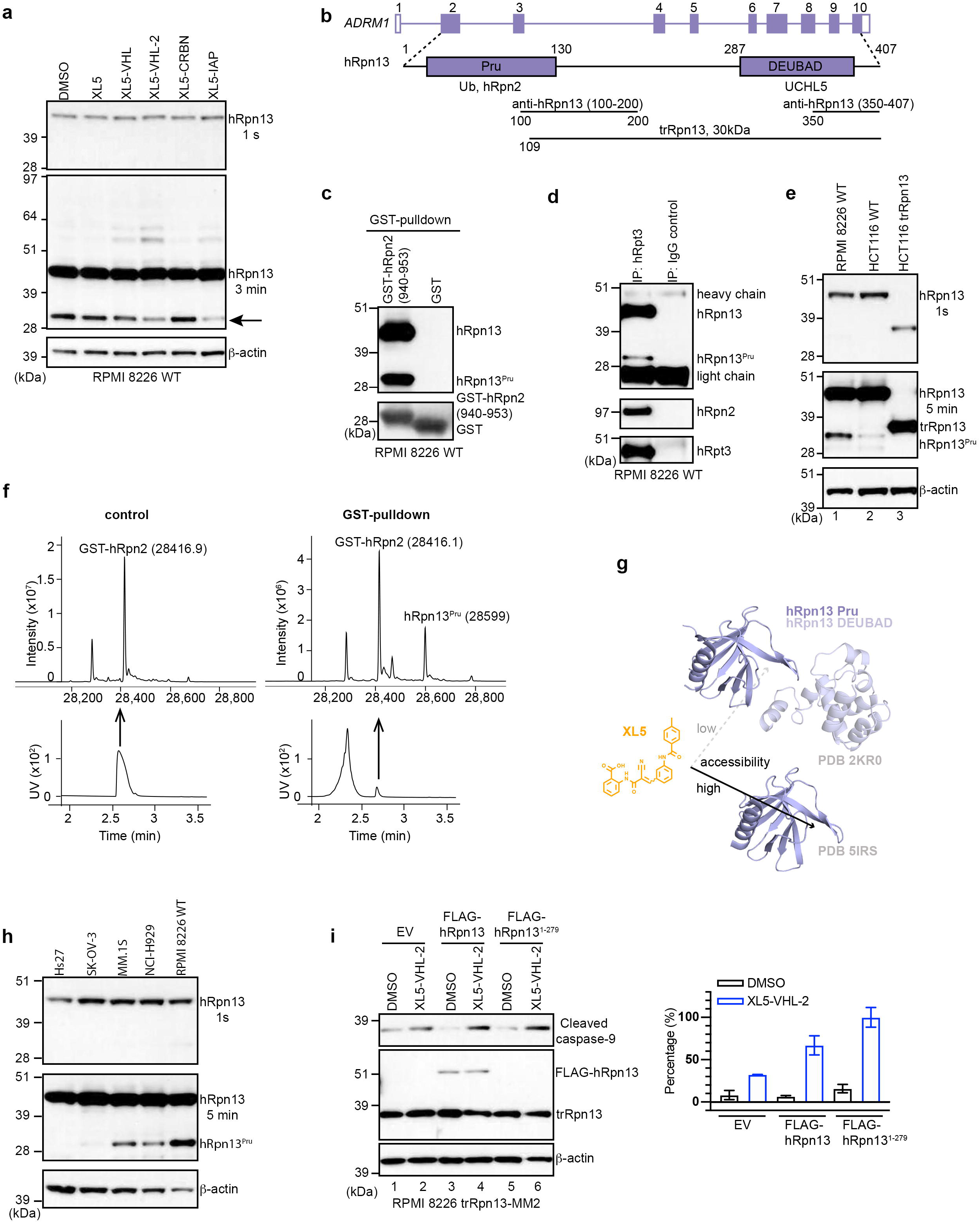
Chemical probing by XL5-PROTACs reveals a DEUBAD-lacking hRpn13 species with cell line dependency. **a,** Immunoblot of whole cell extract from RPMI 8226 WT cells treated for 24 hours with 40 μM XL5, 40 μM XL5-PROTAC or DMSO (vehicle control) detecting hRpn13 (1s or 3 min exposure) or β-actin. **b,** Illustration of hRpn13-encoding *ADRM1* displaying exons, the hRpn13 Pru and DEUBAD domains, the hRpn13 binding sites for ubiquitin (Ub), hRpn2, and UCHL5, the binding epitopes of the two anti-hRpn13 antibodies, and the trRpn13 protein expressed in HCT116 trRpn13 or RPMI 8226 trRpn13-MM1/2 cells. **c**, Immunoblots with antibodies against hRpn13 or GST as indicated from GST pulldowns of RPMI 8226 WT cell lysates. Pulldowns were done with GST-hRpn2 (940-953) or GST (control). **d,** RPMI 8226 WT cells were immunoprecipitated with anti-hRpt3 or IgG (control) antibodies and immunoprobed for hRpn13, hRpn2, and hRpt3 as indicated. **e**, Lysates from RPMI 8226 WT, HCT116 WT, or HCT116 trRpn13 cells were immunoprobed for hRpn13 with 1-second or 5-minute exposure times and β-actin as indicated. **f,** LC-MS analysis of GST-hRpn2 (940-953) (control, left panel) or GST-hRpn2 (940-953)-pulldown sample from lysates of RPMI 8226 WT cells (right panel). The mass spectra (upper panel) were deconvoluted from the UV peak (lower panel) indicated with a black arrow. **g**, Ribbon representation of the structure of hRpn13 Pru domain (solid black line, PDB 5IRS) and full length hRpn13 (dashed grey line, PDB 2KR0) to highlight the greater accessibility to XL5 following loss of the Pru-interacting DEUBAD domain. **h**, Lysates from Hs27, SK-OV-3, MM.1S, NCI-H929, or RPMI 8226 WT cells were immunoprobed for hRpn13 and β-actin as indicated. **i,** Lysates from RPMI 8226 WT and trRpn13-MM2 cells transfected for 48 hours with empty vector (EV) or plasmids expressing FLAG-hRpn13 full length or FLAG-hRpn13^1-279^ proteins were treated for 24 hours with 40 μM XL5-VHL-2 or DMSO (vehicle control) and immunoprobed as indicated with antibodies against hRpn13, cleaved caspase-9, and β-actin (left panel). Quantitation of the levels of cleaved caspase-9 from two independent experiments plotted as mean ± SE as compared to expression of β-actin. Percentage (%) is calculated as the ratio of intensities for cleaved caspase-9 normalized to β-actin (I_cleaved caspase-9_/I_β-actin_)_sample_ divided by that of XL5-VHL-2-treated FLAG-hRpn13^1-279^-expressing cells and multiplied by 100. Plots for DMSO or XL5-VHL-2-treated samples are colored in black or blue, respectively (right panel).

We tested for the presence of hRpn13^Pru^ in HCT116 WT and trRpn13 cells. Lysates from these cell lines and RPMI 8226 WT cells were immunoprobed in parallel with anti-hRpn13 antibodies using β-actin as a loading control. hRpn13 was observed in both WT cell lines and missing in trRpn13 cells (Fig. 5e), as expected. hRpn13^Pru^ was readily observed in RPMI 8226 WT cells, at markedly reduced levels in HCT116 WT cells (Fig. 5e, lane 1 versus lane 2), and absent from HCT116 trRpn13 cells (Fig. 5e, lane 3). The latter finding is consistent with the Exon 2 targeting and results from immunoprobing the two RPMI 8226 trRpn13-MM cell lines (Fig. 4c). HCT116 and RPMI 8226 trRpn13 cells express an hRpn13 protein product that spans M109 to D407 with molecular weight of ∼30 kDa^30^. This truncated hRpn13 protein is slightly larger than hRpn13^Pru^ (Fig. 5e and 4c), which would lack the DEUBAD domain (Fig. 5b). To further characterize hRpn13^Pru^, LC-MS analysis was performed on the GST-hRpn2 pull-down sample. GST-hRpn2 appeared at the expected molecular weight of 28,416.9 Da (Fig. 5f, left panel). From the RPMI 8226 WT cell lysate, a protein was isolated in the GST-hRpn2 pulldown experiment with a mass of 28,599 Da (Fig. 5f, right panel), consistent with hRpn13^Pru^. Assuming no other post-translational modifications, this molecular weight is consistent with an hRpn13 fragment spanning amino acids 1 – 279. That hRpn13^Pru^ is preferentially targeted over full length hRpn13 is consistent with our previous structural data^39^ demonstrating reduced Pru accessibility by interdomain interaction with the DEUBAD (Fig. 5g).

Furthermore, we tested for the presence of hRpn13^Pru^ in non-cancerous skin cell line Hs27 and other cancer cell lines including ovarian cancer cell line SK-OV-3 and multiple myeloma cell lines MM.1S and NCI-H929. The hRpn13 full length protein was upregulated in all cancer cell lines compared to Hs27, whereas only the multiple myeloma cell lines demonstrated upregulated hRpn13^Pru^ (Fig. 5h).

To further interrogate the requirement for XL5-VHL-2 toxicity of full length hRpn13 and/or hRpn13^Pru^, we assessed whether their reintroduction into trRpn13-MM2 cells rescues sensitivity to XL5-VHL-2 treatment. trRpn13-MM2 cells were separately transfected for 48 hours with empty vector (as a control) or expression plasmids for FLAG-hRpn13 full length protein or FLAG-hRpn13^Pru^ (1-279, FLAG-hRpn13^1–279^) and subsequently treated for 24 hours with 40 μM XL5-VHL-2 or DMSO (control). When normalized against β-actin loading control, XL5-VHL-2 treatment induced a small increase in cleaved caspase-9 for the empty vector control (Fig. 5i, left panel, lane 2 versus 1) whereas cells transfected with either full length hRpn13 (Fig. 5i, left panel, lane 4 versus 3) or hRpn13^1-279^ (Fig. 5i, left panel, lane 6 versus 5) indicated a clear induction of caspase-9 cleavage. This effect was reproduced in a second independent experiment and quantified with the resulting plot indicating that either hRpn13 or hRpn13^1-279^ can rescue XL5-VHL-2-driven apoptosis (Fig. 5i, right panel). FLAG-hRpn13 full length protein was detected by anti-hRpn13 antibodies whereas the molecular weight of FLAG-hRpn13^1-279^ at 32 kDa causes it to be indistinguishable from trRpn13 protein (30 kDa, Fig. 5i, left panel). Similarly, any proteolyzed FLAG-hRpn13 to yield FLAG-hRpn13^1-279^ would be indistinguishable from trRpn13.

### hRpn13^Pru^ is depleted in its unmodified form and ubiquitinated following proteasome inhibition

We tested for the presence of a mRNA species corresponding to hRpn13^Pru^ but observed no such mRNA splice variant in our PacBio sequencing data from RPMI 8226 WT cells (Supplementary Fig. 10 and Supplementary Data 1). We next tested whether the proteasome plays a role in generating hRpn13^Pru^. WT RPMI 8226 cells were treated with 100 nM proteasome inhibitor carfilzomib or DMSO (as a control) and immunoprobed for hRpn13 and β-actin (loading control). hRpn13^Pru^ but not full length hRpn13 was reduced following carfilzomib treatment (Fig. 6a, top panel) with corresponding increased abundance for bands between 51 and 64 kDa (Fig. 6a, middle panel) that mirrored those observed with XL5-PROTAC treatment (Fig. 6a, middle panel, 4f and 5a). In addition, a faint upper molecular weight smear was also observed with carfilzomib treatment following hRpn13 immunoprobing. We hypothesized that the upper molecular weight hRpn13 species were ubiquitinated forms stabilized by proteasome inhibition. To assay protein stability, a cycloheximide chase experiment was performed immunoprobing for hRpn13 with β-actin as a loading control. Full length hRpn13 exhibited an apparent half-life of greater than 16 hours whereas that of hRpn13^Pru^ was less than four hours (Fig. 6b). It is possible hRpn13^Pru^ is replenished by cleavage of full length hRpn13 and that its stability is less than indicated by this simple analysis.

**Fig. 6.**
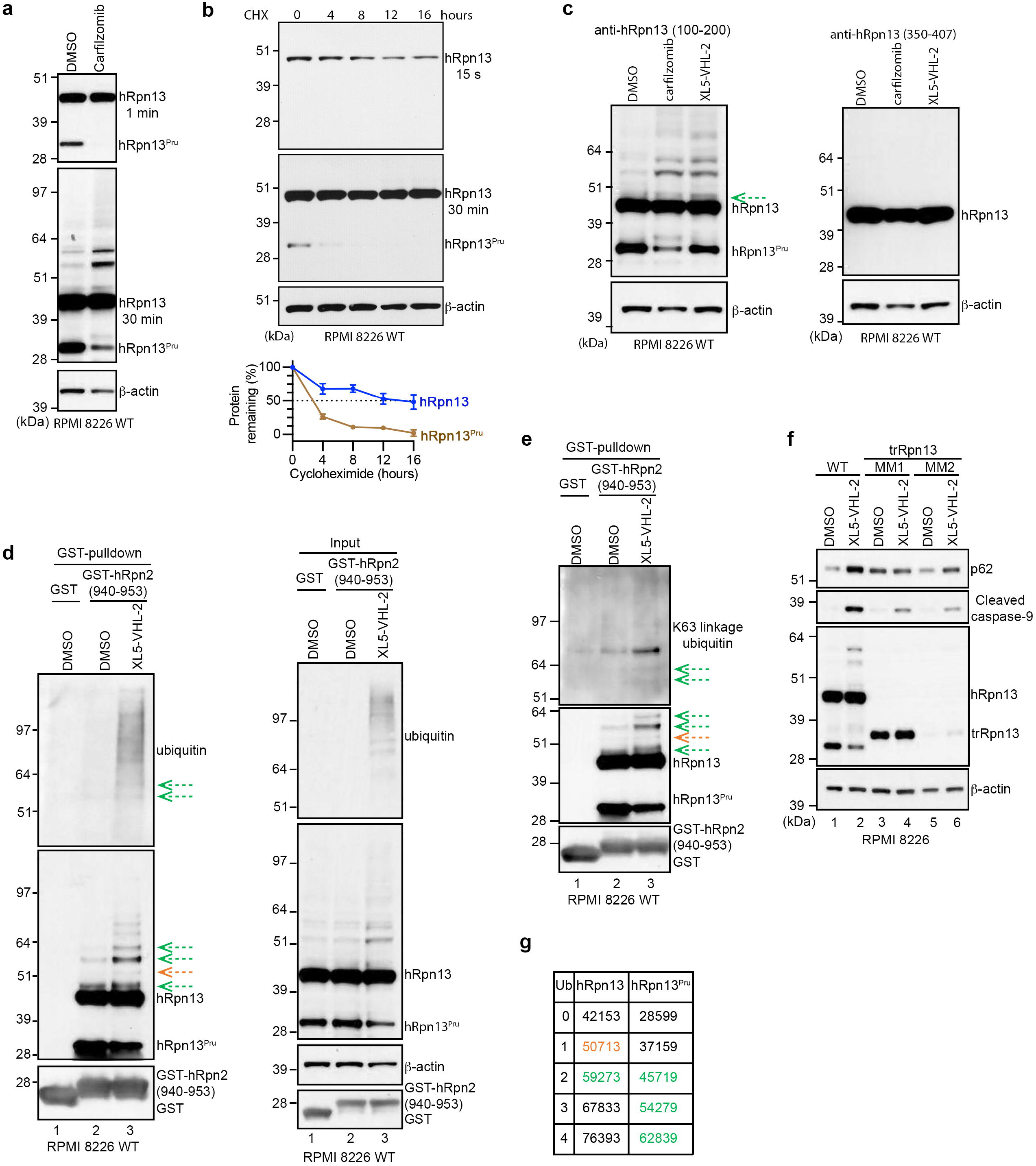
XL5-VHL-2 triggers hRpn13^Pru^ ubiquitination with K63-linkages and increased p62 levels. **a,** Lysates from RPMI 8226 WT cells treated for 24 hours with 100 nM carfilzomib or DMSO vehicle control were immunoprobed for hRpn13 with 1 or 30 minute exposure times and β-actin, as indicated. **b,** Lysates from RPMI 8226 WT cells treated with 50 μg/mL cycloheximide (CHX) for the indicated time points were immunoprobed for hRpn13 and β-actin as a loading control. The immunoblots are representative of three independent experiments. Quantitation of the levels of hRpn13 (from the 15 seconds immunoblot) and hRpn13^Pru^ (from the 30 minutes immunoblot) is reported as mean ± SE in the bottom panel with hRpn13 or hRpn13^Pru^ plotted in blue or brown, respectively. **c,** Lysates from RPMI 8226 WT cells treated with 100 nM carfilzomib, 40 μM XL5-VHL-2 or DMSO (control) for 24 hours were immunoprobed for hRpn13 with anti-hRpn13 antibody recognizing hRpn13 amino acids 100-200 (left panel) or 350-407 (right panel) and β-actin. A faint band above hRpn13 full length protein is marked by a green dashed arrow. **d-e**, Immunoblots with antibodies against ubiquitin, K63 ubiquitin chain linkage (e), hRpn13, GST, or β-actin as indicated of GST pulldowns (left panel, d) and the whole cell lysates (right panel, d) from RPMI 8226 WT cells treated for 24 hours with 40 μM XL5-VHL-2 or DMSO vehicle control. Pulldowns were done with GST-hRpn2 (940-953) or GST (control). Ubiquitinated hRpn13 species are marked by green or orange dashed arrows. **f,** Re-produced experiment as in Fig. 4f with p62 immunoblot included. **g,** Molecular weight of hRpn13 or hRpn13^Pru^ with the conjugation of 0-4 ubiquitin molecules. Ubiquitinated hRpn13 or hRpn13^Pru^ species colored in green or orange match corresponding species indicated with green or orange dashed arrows respectively in c, d, and e.

To provide information on whether the ubiquitinated hRpn13 species correspond to full length hRpn13 or hRpn13^Pru^, we immunoprobed lysates from RPMI 8226 WT cells treated with carfilzomib, XL5-VHL-2, or DMSO with hRpn13 antibodies raised against amino acids 350 – 407 of the DEUBAD domain^46^. Whereas ubiquitinated hRpn13 species were observed as noted above with the Pru/linker domain recognizing antibody, antibodies against the DEUBAD region displayed only unmodified hRpn13 and not the ubiquitinated species (Fig. 6c). As expected from the results of Fig. 5c-d, hRpn13^Pru^ was also not recognized. Altogether, our findings suggest that the ubiquitinated hRpn13 species originate from hRpn13^Pru^ although we cannot preclude the possibility of ubiquitination in the DEUBAD prohibiting recognition of the DEUBAD antibody epitope. Previous studies however have also found hRpn13 to be ubiquitinated upon proteasome inhibition and mapped the ubiquitination sites to Pru domain Lys21 and Lys34^47–49^.

### XL5-VHL-2 induces K63-linked ubiquitination of hRpn13/hRpn13^Pru^ and increases p62 levels

To further characterize the ubiquitinated hRpn13 species generated by XL5-VHL-2 treatment, we used the GST pulldown experiment as applied in Fig. 5c to isolate hRpn13 and its associated proteins through interaction with GST-hRpn2 (940-953). Whole cell lysate from RPMI 8226 WT cells treated with 40 μM XL5-VHL-2 or DMSO vehicle control was incubated with GST-hRpn2 or GST (as a control), mixed separately with glutathione Sepharose 4B resin, which was then washed extensively, and the remaining proteins separated by SDS-PAGE and immunoprobed for ubiquitin, hRpn13 or GST. As expected, GST-hRpn2 pulled down full length hRpn13 and hRpn13^Pru^ as well as the higher molecular weight hRpn13/hRpn13^Pru^ species observed in Fig. 6c (Fig. 6d, left panel). XL5-VHL-2-treated cells exhibited reduced levels of hRpn13^Pru^ and greater abundance of the upper molecular weight bands (Fig. 6d, right panel) as was previously observed (Fig. 4f and 5a). Upper molecular weight bands of >80 kDa were more prominent in the anti-hRpn13 blot of the whole cell lysate than in the GST-hRpn2 pulldown (Fig. 6d, lane 3), suggesting that extensive ubiquitination may interfere with hRpn2 binding.

Immunoprobing for ubiquitin demonstrated the overall presence of ubiquitinated proteins to be increased in the lysates of XL5-VHL-2-treated cells compared to DMSO control (Fig. 6d. right panel). The two distinct bands between 51 and 64 kDa were observed in the anti-ubiquitin immunoblot following pulldown by GST-hRpn2 (Fig. 6d, left panel, lane 3, indicated by dashed arrows), confirming these species to be ubiquitinated hRpn13 or hRpn13^Pru^. Additional ubiquitinated proteins were detected in the GST-hRpn2 pulldown of XL5-VHL-2-treated cell lysates that were not detected by anti-hRpn13 antibodies (Fig. 6d, left panel). These species may represent other ubiquitinated proteins that bind to the hRpn13 Pru and have accumulated by XL5-VHL-2 treatment or alternatively, as proposed above, extensive ubiquitination of hRpn13 may limit antibody detection.

Ubiquitination of proteasome subunits, including hRpn13, induced by starvation or proteasome inhibition is linked to autophagy of the 26S proteasome through a process known as proteophagy^49–52^. Key players in this pathway include K63-linked ubiquitin chains^51^ and the autophagy marker p62, and their interaction is reported to facilitate autophagy-mediated clearance of protein inclusions^53^. To test whether hRpn13 could be involved in proteophagy, we immunoblotted a fresh membrane with the samples included in Fig. 6d (left panel) with a specific antibody against K63-linked ubiquitin chains to assess whether hRpn13 is modified by K63 chains. The two distinct bands between 51 and 64 kDa were detected (Fig. 6e). In addition to these two bands, another band with stronger intensity above 64 kDa was also detected. This band may correspond to hRpn13 or hRpn13^Pru^ modified with a longer ubiquitin chain containing a greater number of K63-linkages. We next assessed the levels of p62 by immunoprobing lysates from RPMI 8226 WT, trRpn13-MM1, or trRpn13-MM2 cells treated with 40 μM XL5-VHL-2 or DMSO (as a control) for p62 in addition to cleaved caspase-9, hRpn13, and β-actin. The effect on p62 level mimicked the differential effect observed for cleaved caspase-9, with a notable increase in p62 for XL5-VHL-2-treated WT cells compared to the two trRpn13-MM cell lines (Fig. 6f and 4f).

Mass spectrometry suggested hRpn13^Pru^ to be of 28,599 Da (Fig. 5f, right panel), as described above. It is mathematically possible that the bands between 51 and 64 kDa correspond to hRpn13^Pru^ at this molecular weight with three and four ubiquitin moieties added respectively (Fig. 6g), as these bands are not recognized by the DEUBAD epitope antibody (Fig. 6c). In addition, a faint band is consistently observed just above where full length hRpn13 protein migrates with the 100 – 200 amino acid epitope that is missing in the DEUBAD-binding epitope (Fig. 6c). This species is consistent in molecular weight with two ubiquitin moieties added to hRpn13^Pru^ (Fig. 6g), but it remains possible that hRpn13 full length protein undergoes a different post-translational modification that causes only slight shifting. Our data suggest that the full length hRpn13 protein is also modified by ubiquitin following XL5-VHL-2 treatment, consistent with the targeting of full length hRpn13 in trRpn13-MM1 (Fig. 4f), as we observe a faint band that best matches monoubiquitination of full length hRpn13 (Fig. 6d-e and 6g, orange) as well as evidence of further hRpn13 ubiquitination (Fig. 6d-g). We note again however that we cannot preclude the possibility of additional post-translational modifications that influence SDS-PAGE migration. Nonetheless, our data collectively indicate a potential role for hRpn13/hRpn13^Pru^ in contributing to proteophagy.

## Discussion

We developed a chemical probe of hRpn13 function that binds with 1.5 μM affinity to the Pru and includes a PROTAC for inducing ubiquitination. XL5 exploits a peripheral cysteine for reversible covalent ligation to hRpn13 with a weak electrophile and non-covalent interactions that mimic those formed at the proteasome by hRpn2. Cysteine-targeting cyanoacrylamide electrophiles form reversible covalent bonds and have been used to inhibit protein kinases with prolonged on-target residence time and higher selectivity^54, 55^ and reversible covalent PROTACs have been developed to degrade kinases with higher selectivity than noncovalent or irreversibly covalent PROTACs^56, 57^. Although XL5 derivatives with modification of the 4-methyl benzamide, benzoic acid or central benzene groups bind hRpn13 with similar or weaker binding affinity than XL5 (Supplementary Table 5 and Supplementary Fig. 7), other modifications may be explored to improve affinity. Beyond the region targeted by XL5, the hRpn2-binding cleft continues where prolines P945, P944, and P942 form myriad interactions. We expect that XL5-VHL-2 and the XL5 general scaffold could be extended to higher affinity by mimicking these interactions (Fig. 3b-c). The differential triggering of apoptosis in multiple myeloma cells based on hRpn13 and hRpn13^Pru^ presence motivates optimization of XL5-VHL-2 for preclinical development.

Between the hRpn13 functional domains is a 157-amino acid linker^39^ of unknown significance. hRpn13^Pru^ appears to extend through this linker region and to accumulate in a ubiquitinated state following proteasome inhibition by carfilzomib (Fig. 6a). It was not noticed in our previous studies with HCT116 cells^30, 35^ due to its low abundance in this cell line (Fig 5e, 1s versus 5 min exposure for hRpn13). The higher expression level in RPMI 8226 cells coupled with the invocation of XL5-PROTACs enabled its identification. We propose that previous publications reporting hRpn13 ubiquitination following proteasome inhibition were likely observing hRpn13^Pru^ ubiquitination^47^. A remaining question is why hRpn13^Pru^ is upregulated in multiple myeloma and how pervasive and frequent it is in other cancer cells. hRpn13^Pru^ appears to be generated by proteasome activity; however, it is not clear whether this is due to a modification of the proteasome or hRpn13 itself. hRpn13^Pru^ harboring the intact Pru but lacking the DEUBAD would be an effective competitor for binding to ubiquitinated substrates and the proteasome, as these intermolecular interactions require displacement of the hRpn13 interdomain interactions^39^. Moreover, hRpn13^Pru^ function would be uncoupled from the UCHL5 deubiquitinase, which hydrolyzes branched ubiquitin chains^58^ and most likely reverses ubiquitination of hRpn13. Additionally, based on the appearance of K63-linked ubiquitin chains (Fig. 6e) in a manner most consistent with hRpn13^Pru^ ubiquitination (Fig. 6g) and on the induction of p62 (Fig. 6f), a compelling model is that acquired ubiquitination of hRpn13^Pru^ contributes to proteophagy, causing reduced proteasome activity. Consequently, these effects would impact the turnover of proteasome substrates in the cell and drive dysregulated cellular proliferation. Furthermore, although XL5-VHL-2 activity is notable on hRpn13^Pru^ in terms of the PROTAC ability to induce target ubiquitination and loss of the unmodified protein (Fig. 4f, 5a, and 6c-g), we cannot exclude the occurrence of these effects on full length hRpn13 (Fig. 4f, 6d-e and 6g), particularly given the effect observed in trRpn13-MM1 cells (Fig. 4f, lane 2 versus 5). It may be that the integrated effects on both hRpn13 and hRpn13^Pru^ drives cellular sensitivity (Fig. 4e) and apoptosis (Fig. 4f).

Altogether our studies have provided new reagents for targeting hRpn13 that uncovered the presence of an hRpn13 species upregulated in multiple myeloma cell lines which are known to be more sensitive to proteasome inhibitors^59^. Specific knockdown of hRpn13^Pru^ without simultaneously targeting full length protein by gene editing or RNAi methods is not feasible. We speculate that full length hRpn13 and hRpn13^Pru^ are both targets of XL5-PROTACs, however, it appears that these compounds preferentially target the more exposed binding surface of hRpn13^Pru^. It is intriguing that carfilzomib treatment appears to yield the same ubiquitinated hRpn13 species (Fig. 6a), suggesting a natural process that is mimicked by PROTAC targeting. Specific targeting of hRpn13^Pru^ preferentially to hRpn13 full length protein may be an effective therapeutic strategy, with less expected toxicity. The mechanism of action for previous hRpn13 targeting compounds failed to be elucidated as interference was not observed for any known hRpn13 activity, including interaction with proteasomes, ubiquitin or UCHL5^20, 27, 31, 32, 35, 60^. This study provides a viable mechanism of action for future investigations of hRpn13 as a therapeutic target.

## Methods

### *In silico* screening

Docking screens were conducted with the ICM-Pro (Molsoft LCC) software^61^ by running up to 1000 parallel processes on 6000 CPUs of the National Institutes of Health Biowulf cluster supercomputer. For the initial screens, the entire hRpn2-binding cleft of hRpn13 was used, including all hRpn13 residues in contact with hRpn2 (940-953), as defined by the NMR and x-ray structures^35, 36^. These amino acids were defined as the targeted binding pocket. Libraries ranged in size from 0.6 to 40 million compounds that were either commercially available (Enamine diversity set, Emolecules, Mcules, Asinex, UORSY, Chembridge, ChemDiv, ChemSpace) or capable of synthesis (Enamine’s diversity REAL database containing 15 million compounds). In total, 63 million compounds were screened. Most of the hits targeted the pocket occupied by the C-terminal end of hRpn2. Enamine’s diversity library of 1.92 million compounds demonstrated the highest hit rate with 5,155 compounds identified in a preliminary fast screen run with a thoroughness value of 1. Hits from the first screens were subjected to more thorough and slow automatic docking with a thoroughness value of 100. 20-30 top compounds from the second round of screens were redocked manually and the best scoring compounds selected for ordering/synthesis and experimental testing.

### Sample preparation

hRpn13 Pru (1-150) or hRpn2 (940-953) was expressed in *E. coli* BL21(DE3) pLysS cells (Invitrogen) as a recombinant protein in frame with an N-terminal histidine tag or glutathione S-transferase respectively followed by a PreScission protease cleavage site. Cells were grown at 37°C to optical density at 600 nm of 0.6 and induced for protein expression by addition of isopropyl-β-D-thiogalactoside (0.4 mM) for 20 hours at 17°C or 4 hours at 37°C. The cells were harvested by centrifugation at 4,550 g for 40 min, lysed by sonication, and cellular debris removed by centrifugation at 31,000 g for 30 min. The supernatant was incubated with Talon Metal Affinity resin (Clontech) for one hour or Glutathione S-sepharose 4B (GE Healthcare Life Sciences) for 3 hours and the resin washed extensively with buffer A (20 mM sodium phosphate, 300 mM NaCl, 10 mM βME, pH 6.5). hRpn13 Pru was eluted from the resin by overnight incubation with 50 units per mL of PreScission protease (GE Healthcare Life Sciences) in buffer B (20 mM sodium phosphate, 50 mM NaCl, 2 mM DTT, pH 6.5) whereas GST-hRpn2 (940-953) was eluted in buffer B containing 20 mM reduced L-glutathione. The eluent was subjected to size exclusion chromatography with a Superdex75 column on an FPLC system for further purification. ^15^N ammonium chloride and ^13^C glucose were used for isotopic labeling.

### NMR experiments

For screening by ^1^H, ^15^N HSQC experiments, small molecule dissolved in DMSO-*d*_6_ was added to 20 μM or 250 μM ^15^N-labeled hRpn13 Pru at a molar excess of 2-fold (for XL5) or 10-fold (for all compounds tested) in NMR buffer (20 mM sodium phosphate, 50 mM NaCl, 2 mM DTT, 10% DMSO-*d*_6_, pH 6.5). All NMR experiments were conducted at 10°C unless indicated to be at 25°C and on Bruker Avance 600, 700, 800 or 850 MHz spectrometers equipped with cryogenically cooled probes. The ^13^C-edited NOESY spectrum was acquired with a 100 ms mixing time on a mixture of 0.4 mM ^13^C-labeled hRpn13 Pru and 0.48 mM unlabeled XL5 in NMR buffer containing 70% ^2^H_2_O. Three ^13^C-half-filtered NOESY experiments were recorded with a 100 ms mixing time on asymmetrically labeled samples dissolved in NMR buffer. One sample contained 0.25 mM ^13^C-labeled hRpn13 Pru mixed with 2-fold molar excess unlabeled XL5; another contained 0.5 mM hRpn13 Pru and 0.5 mM XL5 with the central benzene ring ^13^C-labeled (XL5 ^13^C_6_-CB); and a third contained 0.4 mM hRpn13 Pru and 0.4 mM XL5 with the benzoic acid ring ^13^C-labeled (XL5 ^13^C_6_-BA) dissolved in NMR buffer containing 70% ^2^H_2_O. An ^15^N-dispersed NOESY spectrum was acquired with a 120 ms mixing time on 0.25 mM ^15^N-labeled hRpn13 Pru mixed with 2-fold molar excess unlabeled XL5 dissolved in NMR buffer. The ^1^H, ^13^C HMQC experiments were acquired on 0.5 mM XL5-^13^C_6_-CB in NMR buffer with and without DTT as well as mixed with equimolar unlabeled hRpn13 Pru; a control experiment with only 0.5 mM hRpn13 Pru was also recorded in NMR buffer to assign natural abundance signals of hRpn13. 2D ^13^C-edited HCCH-TOCSY (12 ms mixing time), NOESY (500 ms mixing time), or ^1^H, ^13^C HMQC spectra were recorded on 10 mM XL5-^13^C_6_-BA in DMSO-*d*_6_ at 25°C, and ^1^H, ^13^C HMQC spectra were recorded in NMR buffer on 0.1 mM XL5-^13^C_6_-BA with increasing molar ratio of unlabeled hRpn13 Pru, including at 1:0, 1:0.5, 1:1, 1:2, and 1:4. Data were processed by NMRPipe^62^ and visualized with XEASY^63^.

### Chemical shift assignments

Chemical shift assignments for hRpn13 were aided by a previous study^35^ and confirmed by NOESY experiments; namely, an ^15^N-dispersed NOESY (120 ms mixing time) experiment recorded in NMR buffer on 0.25 mM ^15^N hRpn13 Pru mixed with 2-fold molar excess XL5 or a ^13^C-edited NOESY (100 ms mixing time) experiment recorded on a mixture of 0.48 mM unlabeled XL5 and 0.4 mM ^13^C labeled hRpn13 Pru dissolved in NMR buffer with 70% ^2^H_2_O.

To aid in the chemical shift assignment of XL5, we selectively ^13^C-labeled either the benzoic acid aromatic ring (Supplementary Fig. 2a, top panel) or the central benzene ring (Supplementary Fig. 2a, bottom panel); we refer to these samples as XL5-^13^C_6_-BA and XL5-^13^C_6_-CB respectively. H15, H16, H17 and H18 from XL5 were assigned by using ^13^C-edited 2D HCCH-TOCSY, 2D NOESY and HMQC spectra recorded on 10 mM XL5-^13^C_6_-BA in DMSO-*d*_6_ (Supplementary Fig. 2b-c). These assignments could be transferred for XL5 dissolved in NMR buffer although shifting and splitting was observed due to the presence of 2 mM DTT (Supplementary Fig. 2d, left most spectrum). Addition of unlabeled hRpn13 Pru caused shifting for XL5 H17 and H18, as well as the H15 and H16 signals to attenuate (Supplementary Fig. 2d). Without DTT, the four expected signals for H9, H10, H11 and H12 appeared in the spectrum recorded on XL5-^13^C_6_-CB; however, inclusion of DTT in the NMR buffer caused multiple new signals to appear (Supplementary Fig. 2e, middle panel versus left panel), as was observed for the XL5 benzoic acid group (Supplementary Fig. 2c-d). Addition of hRpn13 Pru caused all XL5-^13^C_6_-CB signals present in the ^1^H, ^13^C HMQC spectrum to disappear with the exception of one weak signal (Supplementary Fig. 2e, right panel); this resonance was assigned to H12 by an NOE interaction to H8 of XL5 that was observed in a ^1^H, ^13^C half-filtered NOESY experiment recorded on 0.5 mM XL5-^13^C_6_-CB mixed with equimolar unlabeled hRpn13 Pru (Supplementary Fig. 2f).

### Structure determination

Distance, dihedral angle and hydrogen bond restraints were generated from the unligated hRpn13 Pru crystal structure (PDB 5IRS)^9^ with the exception of amino acids at the binding interface, including M31, L33, V38, T39, V85 V93 and F106, for which restraints from the spectra recorded on XL5-ligated hRpn13 were used exclusively to allow for rearrangements due to XL5 binding. These restraints were combined with 23 NOE-derived distance restraints between hRpn13 and XL5 (Fig. 2a-b, Table 1 and Supplementary Table 3) to calculate the XL5-ligated hRpn13 Pru structure. The calculations were done by using simulated annealing algorithms in XPLOR-NIH 2.50 (http://nmr.cit.nih.gov/xplor-nih/)^64^. An initial set of topology and parameter files for the ligand were generated by PRODRG^65^ and corrected to require the angles in the planar 6-membered rings to sum to 360°. XL5 was covalently bonded to the hRpn13 C88 sulfur of PDB 5IRS (as displayed in Fig. 2a) with chirality at XL5 C15 and C16 of S, S (SS), R, R (RR), S, R (SR) or R, S (RS) stereochemistry. Each stereoisomer was used as a starting structure for iterative simulated annealing to generate 200 initial structures, from which twenty were chosen based on criteria of no NOE, dihedral or torsion angle violation and lowest energy. The structures were then clustered into converged sets (Supplementary Fig. 5) and evaluated based on adherence to differential NMR data such that distances were closer for interacting protons with stronger NOEs. The only structures that fit all of the NMR data were those of SS stereochemistry and in the main cluster 1 which contained seventeen of the twenty calculated SS structures. This cluster places XL5 H17 closer to hRpn13 L33 Hγ than XL5 H18 and XL5 H18 closer to a hRpn13 V38 methyl group than XL5 H17 and H15. These differential interactions are indicated by the stronger NOEs observed between XL5 H17 or H18 with hRpn13 L33 Hγ or V38 methyl group respectively (Fig. 2a-b) and not preserved in cluster 2 (Supplementary Fig. 5). The calculated RS and SR structures formed four clusters whereas the RR structures formed six clusters; however, these clusters failed to fit the NMR data, such as the directing of RS cluster 1 or SR cluster 3 XL5 H13 away from hRpn13 V85 (Fig. 2a and Supplementary Fig. 5) or yielding equivalent interactions for XL5 H19, H15, H17 or H18 with the observed hRpn13 V38 methyl group as occurs in RS cluster 2-4, SR cluster 1, 2 and 4, and RR cluster 1-4 (Fig. 2a-b and Supplementary Fig. 5). Similarly, the closer proximity in RR cluster 5 and cluster 6 of the hRpn13 V85 methyl groups to XL5 H19 than XL5 H13 is not supported by NMR data (Fig. 2a and Supplementary Fig. 5). Altogether, our structure calculations best support XL5 binding to hRpn13 with SS chirality for XL5 C15 and C16; however, we cannot preclude the possibility of small populations existing with XL5 SR, RS or RR chirality.

A weak hydrogen bond between the hRpn13 S90 sidechain hydroxy group and XL5 cyanide group was found in eight of the SS cluster 1 structures. Therefore, this hydrogen bond was included as an additional distance restraint (Table 1) and a new iteration of SS structure calculations was performed to yield 20 final lowest energy structures without hRpn13 distance or dihedral angle violations greater than 0.5 Å or 5° respectively and no torsion angle violations. This final set of 20 structures was selected for visualization and statistical analyses. Structure evaluation was performed with the program PROCHECK-NMR^66^; the percentage of residues in the most favored, additionally allowed, generously allowed and disallowed regions was 94.3, 5.7, 0.1 and 0.0, respectively. Visualization was performed with MOLMOL^67^ or PyMOL (PyMOL Molecular Graphics System, https://www.pymol.org/2/).

### DSF

DSF experiments were performed on a Prometheus NT.48 instrument (NanoTemper Technologies, Germany) at 20°C. 40 μM compound was added to equal volume of 2 μM hRpn13 Pru in buffer C (20 mM sodium phosphate, 50 mM NaCl, 10% DMSO, pH 6.5). For Fig. 1a, 2 μM hRpn13 Pru was added to equal volume of serially diluted XL5 or RA190 in buffer C. Each sample was loaded into three capillaries of High Sensitivity grade (NanoTemper, cat # PR-C006) and the emission of intrinsic tryptophan fluorescence at 350 nm was monitored.

### ITC experiments

ITC experiments were performed at 25°C on a MicroCal iTC200 system (Malvern, PA, USA). hRpn13 Pru, XL5, XL5 derivative, or RA190 were prepared in buffer C. One aliquot of 0.5 μL followed by 17 or 18 aliquots of 2.1 μL of 200 μM hRpn13 Pru was injected at 750 r.p.m. into a calorimeter cell (volume 200.7 ml) that contained 20 μM XL5, XL5 derivative, or RA190. Blank experiments were performed by replacing XL5, XL5 derivative, or RA190 with buffer in the cell and the resulting data subtracted from the experimental data during analyses. The integrated interaction heat values were normalized as a function of protein concentration and the data were fit with MicroCal Origin 7.0 -based software implementing the “One Set of Sites” model to yield binding affinity *K*_a_ (1/*K*_d_), stoichiometry, and other thermodynamic parameters.

### LC-MS experiments

LC-MS experiments were performed on a 6520 Accurate-Mass Q-TOF LC/MS system equipped with a dual electro-spray source, operated in the positive-ion mode. Samples included 2 μM hRpn13 Pru incubated for 2 hours at 4°C with 10-fold molar excess XL5 in buffer C containing 0.2% DMSO as well as 2 mM reduced L-glutathione incubated for 2 hours at 4°C with 40 μM XL5 or RA190 in buffer C containing 0.4% DMSO. Acetonitrile was added to all samples to a final concentration of 10%. Data acquisition and analysis were performed by Mass Hunter Workstation (version B.06.01). For data analysis and deconvolution of mass spectra, Mass Hunter Qualitative Analysis software (version B.07.00) with Bioconfirm Workflow was used.

To check for reactivity of XL5 in mouse serum, 0.2 µM XL5 was mixed with mouse serum (BioIVT) and aliquots of the spiked mixture left at room temperature for 0, 4, 8 and 24 hours. For each time point, six samples were extracted using 75% acetonitrile and 0.075% formic acid. The supernatant was transferred to polypropylene injection vials for LC-MS analysis. LC-MS was performed with a TSQ Quantiva triple quadrupole mass spectrometer (Thermo Fisher Scientific) operating in selected reaction monitoring mode with positive electrospray ionization and with a Shimadzu 20AC-XR system using a 2.1 x 50 mm, 2.7 µm Waters Cortecs C18 column.

### Acquisition of compounds

XL1 (CAS:860-22-0) was purchased from Sigma-Aldrich; XL2-XL22 (Enamine ID listed in the Supplementary Table 1) and XL23 (Enamine ID Z44395249) were ordered from Enamine; XL5-^13^C_6_-BA, XL24, XL28 and XL29 were obtained by customized synthesis from Enamine; XL5-^13^C_6_-CB, XL25, XL26, XL27, XL30, XL31, XL32, XL33, XL5-VHL, XL5-VHL-2, XL5-CRBN, XL5-IAP, VHL-Ac, IAP-Bz were synthesized according to the reported literature procedures^41, 68^ and described in the Supplementary Note.

### Generation of trRpn13 RPMI-8226 cell lines

trRpn13-MM1 and trRpn13-MM2 cells were generated using the CRISPR/Cas9 system. Six candidate sgRNAs were designed by using the sgRNA Scorer 2.0 web tool^69^ and subsequently tested for activity in 293T cells with a previously described approach^70^ (Supplementary Table 6). Candidates 2288 and 2290 were identified to be the most potent and were used for further experiments. Three different combinations of Cas9/sgRNA were generated and used: Cas9/2288, Cas9/2290, and Cas9/2288/2290. For each condition, 4 μg of *in vitro* transcribed guide RNA was complexed with 10 μg of purified recombinant Cas9 protein and electroporated into 200,000 cells by the default RPMI 8226 settings of the Lonza 4D Nucleofector system. Seventy-two hours later, cells were stained with propidium iodide (PI) for viability and single cell sorted into 96-well plates. For each combination, 2 – 96-well plates were sorted and allowed to grow for ∼6 weeks. Upon sufficient growth, genomic DNA was isolated from 74 viable clones by a solution-based DNA extraction method^71^ and screened with targeted Illumina sequencing^70^. Eleven clonal populations were identified to have the putative genetic disruption of which only two were able to survive long term culture (trRpn13-MM1, trRpn13-MM2). One additional clone, which did not have any editing, was retained and served as an experimental control. Genomic DNA from control cells and trRpn13 MM cells were extracted by QIAamp^→^ DNA Mini Kit (51304; Qiagen). DNA editing status was validated by performing TOPO blunt cloning of PCR amplicons encompassing the guide RNA target sites and Sanger sequencing. Primers for PCR product generation are provided in Supplementary Table 7.

### Cell culture and antibodies

The HCT116 WT (ATCC^®^CCL-247™), RPMI 8226 (ATCC^®^ CCL-155™), Hs27 (ATCC^®^ CRL-1634™), SK-OV-3 (ATCC^®^HTB-77), MM.1S (ATCC^®^ CRL-2974™) and NCI-H929 (ATCC^®^ CRL-9608™) cell lines were purchased from the American Tissue Culture Collection; HCT116 trRpn13 cells were generated and described as part of a previous study^30^. HCT116 and SK-OV-3 cell lines were grown in McCoy’s 5A modified media (Thermo Fisher Scientific 16600082); RPMI 8226 cell lines, MM.1S and NCI-H929 cell lines were grown in RPMI-1640 media (ATCC® 30-2001™); the Hs27 cell line was grown in DMEM media (Thermo Fisher Scientific, 10569010). In all cases, the media was supplemented with 10% fetal bovine serum (Atlanta Biologicals) and growth occurred in a 37°C humidified atmosphere of 5% CO_2_. 0.05 mM βME was added to the media for HCI-H929 cells. Antibodies (dilutions) used in this study include anti-hRpn13 (100-200) (Abcam ab157185, 1:5,000), anti-hRpn13 (350-407) (Abcam ab157218, 1:5,000), anti-hRpn2 (Abcam ab2941, 1:1,000), anti-hRpt3 (Abcam ab140515, 1:1,000), anti-UCHL5 (Abcam ab133508, 1:2,000), anti-β-actin (Cell Signaling Technology 4970s or 3700s, 1:3,000, 1:5000 or 1:10,000), anti-cleaved caspase-9 (Cell Signaling, 52873s, 1:500), anti-ubiquitin (P4D1) (Cell Signaling, 3936s, 1/1000), anti-K63 ubiquitin chain linkage (Cell Signaling, 5621s, 1/1000), anti-GST (Cell Signaling, 2625s, 1:10,000), anti-mouse (Sigma-Aldrich, A9917, 1:3,000 or 1:4,000), anti-rabbit (Life Technologies, A16110, 1:5,000, 1:10,000 or 1:20,000) and anti-native rabbit (Sigma-Aldrich, R3155, 1:1000) antibodies.

### MTT assay

HCT116 WT or trRpn13 cells were seeded at 4,000 cells/well whereas RPMI 8226 WT, trRpn13-MM1 or trRpn13-MM2 cells were seeded at 8,000 cells/well with RPMI 1640 medium (no phenol red, Thermo Fisher Scientific 11835030) containing 2% fetal bovine serum in 96-well plates. After 24 hours, cells were treated with 0.4% DMSO (as a control) and this concentration was maintained with XL5, XL5-PROTACs XL5-VHL, XL5-VHL-2, XL5-CRBN, XL5-IAP, E3 ligand VHL-Ac, thalidomide (Selleckchem, catalog NO. S1193), or IAP-Bz at 10 µM, 20 µM, 30 µM or 40 µM concentration. RPMI 8226 WT, trRpn13-MM1 or trRpn13-MM2 were treated similarly but with XL5-VHL-2 at 2.5 µM or 5 µM concentration. Each condition was performed in sextuplicate. After 48 hours, 0.35 mg/mL MTT was added for 4 hours of incubation. Stop solution (40% DMF, 10% SDS (W/V), 25 mM HCl, 2.5% acetic acid in H_2_O) was added to the cells and incubated overnight. Absorbance at 570 nm was measured by using CLARIOstar (BMG LABTECH).

### XL5 treatment

Two million RPMI 8226 WT, trRpn13-MM1 or trRpn13-MM2 cells were seeded in T75 flask. After 48 hours, the cells were treated with 40 μM XL5, 40 μM XL5-PROTAC, 100 nM carfilzomib or 0.8% DMSO (as a control) for 24 hours, as indicated.

### Cycloheximide

After 24 hours of plating, at time point 0, RPMI 8226 WT cells were treated with cycloheximide (50μg/mL) for 4, 8, 12 and 16 hours. At each time point, cells were harvested, washed with PBS, then flash frozen in liquid nitrogen before storing at −80°C until processing for immunoprobing. Protein expression levels were quantitated by using Image Studio (version 2.5.2, Licor) and normalized to β-actin.

### Immunoblotting

HCT116 WT, HCT116 trRpn13, RPMI 8226 WT, RPMI 8226 trRpn13-MM1, RPMI 8226 trRpn13-MM2, Hs27, SK-OV-3 or NCI-H929 cells were collected and washed with PBS followed by flash freezing in liquid nitrogen and storage at −80°C. Cells were lysed in 1% Triton-TBS lysis buffer (50 mM Tris-HCl, pH 7.5, 150 mM NaCl, 1mM PMSF) supplemented with protease inhibitor cocktail (Roche). Total protein concentration was determined by bicinchoninic acid (Pierce). Protein lysates were prepared in 1x LDS (ThermoFisher, NP0007) buffer with 100 mM DTT and heating at 70°C for 10 min, loaded onto 4–12% Bis-Tris polyacrylamide gels (Life Technologies), subjected to SDS–PAGE and transferred to Invitrolon polyvinylidene difluoride membranes (Life Technologies). The membranes were blocked in Tris-buffered saline with 0.1% Tween-20 (TBST) supplemented with 5% skim milk or 5% BSA, incubated with primary antibody, washed in TBST, incubated with secondary antibodies and washed extensively in TBST. Pierce^TM^ ECL Western Blotting Substrate (32106; Thermo Fisher Scientific) or Amersham^TM^ ECL^TM^ Primer Western Blotting Detection Reagent (cytiva) was used for antibody signal detection.

### Plasmids for transfection

Plasmids expressing FLAG-tagged hRpn13 or hRpn13^1-279^ were generated commercially (GenScript) by inserting synthesized coding sequence for full length hRpn13 (NM_007002.3) or for residues 1–279 between the BamHI and HindIII restriction sites of pCMV-3Tag-1a (Agilent Technologies, 240295). Unmodified pCMV-3Tag-1a was used as empty vector (EV) control.

### Transfection

RPMI 8226 (WT and trRpn13-MM2) cells (2.5 x 10^5^) were reverse transfected with 1 μg empty vector (EV), 2.5 μg FLAG-hRpn13-expressing plasmid or 5 μg FLAG-hRpn13^1-279^-expressing plasmid by using lipofectamine 3000 (Thermo Fisher Scientific, L3000015) according to the manufacturer’s instructions. After 48 hours of transfection, cells were treated with 40 μM XL5-VHL-2 or DMSO (vehicle control) for 24 hours before harvesting cells.

### Immunoprecipitation

RPMI 8226 cell lysates (1mg) were incubated with anti-hRpt3 (Abcam ab140515, 1:100) or IgG (rabbit) antibodies overnight at 4°C and then incubated for an additional 3 hours at 4°C with 50 μL Dynabeads^TM^ protein G (Life Technologies, 10004D). Following three washes with 1% Triton-TBS lysis buffer, proteins bound to the Dynabeads^TM^ protein G were eluted by using 2x LDS with 100 mM DTT and analyzed by immunoblotting.

### GST-pulldown assay

RPMI 8226 cell lysates (1-2 mg) were incubated with 2 nmol GST or purified GST-hRpn2 (940-953) overnight at 4°C and then incubated for an additional 3 hours at 4°C with 25 μL pre-washed glutathione Sepharose 4B resin (cytiva). Following three washes with 1% Triton-TBS lysis buffer, proteins bound to the glutathione Sepharose 4B resin were eluted by using 2x LDS with 100 mM DTT and analyzed by immunoblotting or eluted in 50 μL buffer B containing 20 mM reduced L-glutathione for LC-MS analysis as described in LC-MS experiments.

### RNA PacBio Sequencing and Illumina sequencing

Total RNA sample from RPMI 8226 WT, trRpn13-MM1 or trRpn13-MM2 cells were extracted by using the RNeasy Plus minikit (74134; Qiagen). RNA PacBio sequencing was performed on each sample, and Illumina sequencing was additionally performed on the RPMI 8226 WT sample. RNA was used for full-length transcript sequencing by the PacBio Sequel II system (Pacific Biosciences, CA, USA). Iso-Seq libraries were prepared following the Iso-Seq™ Express Template Preparation protocol (PN 101-763-800, Pacific Biosciences, CA) and size selected using ProNex beads (Promega) with a bead-to-DNA ratio of 0.95 to incorporate transcripts < 2kb. Each library was then sequenced by a SMRT Cell 8M with the PacBio Sequel II platform according to the manufacturer’s instructions. The library preparation used Illumina TruSeq Stranded mRNA LT kit (RS-122-2101). The Poly-A containing mRNA molecules are purified using poly-T oligo attached magnetic beads. Following purification, the mRNA is fragmented into small pieces and the cleaved RNA fragments are copied into first strand cDNA using reverse transcriptase and random primers, followed by second strand cDNA synthesis using DNA Polymerase I and RNase H. The resulting double-strand cDNA is used as the input to a standard Illumina library prep with end-repair, adapter ligation and PCR amplification being performed to produce a sequencing ready library. The final purified product is then quantitated by qPCR before cluster generation and sequencing. The libraries were run on the Illumina NextSeq instrument using NextSeq High v2.1 kit and run as 2×76bp paired-end sequencing run.

### Illumina-short Reads Transcriptomic Sequencing Analysis

The HiSeq Real Time Analysis software (RTA 2.11.3) was used for processing raw data files, the Illumina bcl2fastq2.17 was used to demultiplex and convert binary base calls and qualities to fastq format. The sequencing reads were trimmed adapters and low quality bases using Cutadapt (version 1.18), the trimmed reads were mapped to human reference genome (hg38) and Gencode annotation GENCODE v30 using STAR (version 2.7.0f) with two-pass alignment option. RSEM (version 1.3.1) was used for gene and transcript quantification.

### PacBio Iso-seq Analysis

Raw subreads were converted into HiFi circular consensus sequences (CCS). The CCS reads were processed using the isoform sequencing (IsoSeq v3) pipeline by demultiplexing the barcodes and removing primers. Additional refine steps included trimming polyA tails and removing concatemers to generate Full Length Non-Concatemer (FLNC) reads. Iterative clustering was performed to obtain consensus isoforms, and the full-length (FL) consensus sequences. The high-quality full-length transcripts were classified based on a post-correction accuracy criterion of above 99%. The FL consensus sequences were mapped to the reference genome by using the minimap2 software. Transcript annotation is done using squanti3 software and Illumina short-read RSEM gene expression data was integrated with PacBio Iso-seq transcript for quantification.

### Statistical information

The violations and deviations from idealized geometry in Table 1 were obtained by XPLOR-NIH and average pairwise root-mean-square deviation were calculated by MOLMOL.

Mean values, standard deviation and standard error were calculated by using Microsoft Excel. The values for n represent replicates of biochemical assays displayed in Fig. 1a, 1e, 4b, 4e, 5i (right panel), 6b (bottom panel), Supplementary Fig. 3c and Supplementary Table 1. For each figure or table, the number of replicates is indicated in the figure or table legend.

The *K*_d_ values in Fig. 1d, Supplementary Fig. 1c, 7 and Supplementary Table 5 were generated by fitting ITC data to a “One Set of Sites” binding model with the Origin software. IC_50_ values in Fig. 4b (right panel) were analyzed from the data in Fig. 4b (left panel) by using the equation [Inhibitor] vs. normalized response] in GraphPad Prism8. Chemical shift perturbation (CSP) analysis was done by comparing ^1^H, ^15^N HSQC experiments recorded on ^15^N-labeled hRpn13 Pru alone and with 2-fold molar excess unlabeled XL5. CSP values were calculated according to Eq 1^72^, where Δδ_N_ and Δδ_H_ symbolize change in amide and proton signal, respectively, and a threshold of one standard deviation above average was used for the plot (Supplementary Fig. 1b).

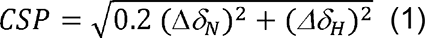

Mass Hunter Qualitative Analysis software (version B.07.00) with Bioconfirm Workflow was used to deconvolute mass spectra and integrate UV spectra in Fig. 2c, Fig. 2d, Fig. 5f and Supplementary Fig. 3a and 3b.

### Data Availability

The structural coordinates and chemical shift data for XL5-ligated hRpn13 Pru has been deposited into the Protein Data Bank (PDB) and Biological Magnetic Resonance Data Bank (BMRB) with accession codes PDB: 7KXI and BMRB: 30824.

## Acknowledgements

This work was supported by the Intramural Research Program of the CCR, NCI, NIH (1 ZIA BC011490 and 1 ZIA BC011627). We gratefully thank Janusz Koscielniak for his maintenance of the NMR spectrometers, Karen Stefanisko for her suggestions regarding the MTT assay, Burchelle Blackman (NHLBI) for obtaining high resolution mass spectrometry data, and Gwen Buel for critical reading of the manuscript. This work utilized the computational resources of the NIH HPC Biowulf cluster (http://hpc.nih.gov).

## Author contributions

K.J.W. and X.L. conceived of the project. X.L. performed the NMR and DSF experiments and solved the XL5-ligated hRpn13 structure. X.L. performed all of the cell biology experiments except for the cycloheximide chase and rescue experiments, which were done by V.O.A.; N.I.T. performed the *in silico* screening. ITC was done by X.L. and S.G.T.; LC-MS experiments except for in mouse serum were acquired by M.D. and analyzed by X.L. and M.D.; K.C.C. performed LC-MS for samples with mouse serum, which were designed by T. A. and K.C.C.; V.S. synthesized XL5 with the central benzene ^13^C labeled, XL25-XL27, XL30-XL33 and XL5 PROTACs, which were designed by R.S., K.J.W., and V.S.; X.L. generated all of NMR samples except for one generated by X.C.; C.D.S. generated modified topology and parameter files for XL5. X.L. and H.M. performed NMR experiments on free XL5 with the benzoic acid group ^13^C labeled; X.L. performed all other NMR experiments. RNA samples and gDNA samples for sequencing were prepared by X.L. CRISPR-edited RPMI 8226 trRpn13 MM cell lines were designed by R.C., generated by C.N.E. and validated by J.C.K. with supervision by R.C.; RNA sequencing was conducted by C.F. and analyzed by S.C. with supervision by Y.Z. and B.T.; K.J.W. and X.L. wrote the manuscript with contribution from all authors.

