## Supplementary Information for "Structure-guided bifunctional molecules hit a DEUBAD-lacking hRpn13 species upregulated in multiple myeloma"

**Abstract:** Proteasome substrate receptor hRpn13 is a promising anti-cancer target. By integrated *in silico* and biophysical screening, we identified a chemical scaffold that binds hRpn13 with non-covalent interactions that mimic the proteasome and a weak electrophile for Michael addition. hRpn13 Pru domain binds proteasomes and ubiquitin whereas its DEUBAD domain binds deubiquitinating enzyme UCHL5. NMR revealed lead compound XL5 to interdigitate into a hydrophobic pocket created by lateral movement of a Pru β-hairpin with an exposed end for Proteolysis Targeting Chimeras (PROTACs). Implementing XL5-PROTACs as chemical probes identified a DEUBAD-lacking hRpn13 species (hRpn13$^{Pru}$) present naturally with cell type-dependent abundance. XL5-PROTACs preferentially target hRpn13$^{Pru}$, causing its ubiquitination. Gene-editing and rescue experiments established hRpn13 requirement for XL5-PROTAC-triggered apoptosis and increased p62 levels. These data establish hRpn13 as an anti-cancer target for multiple myeloma and introduce an hRpn13-targeting scaffold that can be optimized for preclinical trials against hRpn13$^{Pru}$-producing cancer types.

**Supplementary Table 1. Twenty-two lead compounds identified from an *in silico* screen as putative hRpn13 binders.**

| Nomenclature | ID | Validation Method | $\lambda_{350}$ (normalized) |
|---|---|---|---|
| DMSO | D1435 or DLM-34-10X0.75 | NMR, DSF, ITC | 100 ± 3.0 |
| XL1 | CAS:860-22-0 | DSF | 81.5 ± 2.9 |
| XL2 | Z119018758 | DSF | 86.1 ± 0.3 |
| XL3 | Z231949652 | DSF | 67.9 ± 3.0 |
| XL4 | Z17870320 | DSF | 64.8 ± 1.1 |
| XL5 | Z44395247 | NMR, DSF, ITC | 47.2 ± 0.5 |
| XL6 | Z45668530 | NMR, DSF | 78.4 ± 1.1 |
| XL7 | Z2301703555 | NMR, DSF | 80.2 ± 1.5 |
| XL8 | Z57354452 | DSF | 76.2 ± 0.6 |
| XL9 | Z199467950 | DSF | 83.1 ± 1.8 |
| XL10 | Z56774971 | DSF | 82.1 ± 1.6 |
| XL11 | Z211851080 | DSF | 85.9 ± 0.4 |
| XL12 | Z87597408 | DSF | 84.6 ± 0.8 |
| XL13 | Z146687966 | DSF | 87.7 ± 1.7 |
| XL14 | Z96452288 | DSF | 84.4 ± 5.9 |
| XL15 | Z217127446 | DSF | 75.0 ± 0.8 |
| XL16 | Z2154430633 | NMR, DSF | 81.1 ± 1.6 |
| XL17 | Z1082901996 | NMR, DSF | 76.9 ± 1.2 |
| XL18 | Z1917789752 | NMR, DSF | 78.5 ± 0.8 |
| XL19 | Z2154430997 | NMR, DSF | 65.1 ± 5.1 |
| XL20 | Z2910888840 | NMR, DSF | 86.2 ± 5.0 |
| XL21 | Z1262429908 | NMR | N/A |
| XL22 | Z3039488982 | NMR | N/A |
| RA190 | M60163-2s | DSF, ITC | 76.7 ± 2.1 |

Nomenclature, Enamine ID (XL2-XL22), CAS number (XL1), Xcessbio ID (RA190, positive control), Sigma-Aldrich or Cambridge Isotope Laboratories, Inc ID (DMSO, negative control), biophysical method used to screen compounds, and normalized intrinsic tryptophan fluorescence emission is listed in column 1, 2, 3, and 4 respectively. Emission of intrinsic tryptophan fluorescence was measured at 350 nm in triplicate for 1 μM hRpn13 Pru or with addition of 20-fold molar excess of listed compound. The average fluorescence intensity at 350 nm for each sample was normalized to hRpn13 Pru with DMSO addition and is presented with the standard deviation from the mean in column 4. N/A, not available.

**Supplementary Table 2. Chemical shift assignments for hRpn13-bound XL5.**

| XL5 | chemical shift (ppm) |
|---|---|
| $CH_3$ | 2.376 |
| H4, H7 | 7.360 |
| H5, H6 | 7.688 |
| H8 | 10.274 |
| H12 | 7.258 |
| H13 | 3.648 |
| H15 | 7.934 |
| H16 | 6.802 |
| H17 | 7.054 |
| H18 | 7.866 |
| H19 | 4.672 |

**Supplementary Table 3. NOE interactions detected between hRpn13 and XL5.**

| hRpn13 | XL5 |
|---|---|
| M31 $CH_3$ | H15, H19 |
| L33 H$\beta$ | H17, H18 |
| L33 H$\gamma$ | H17, H18 |
| L33 $CH_3$ | H16, H17, H18 |
| V38 $CH_3$ | H17, H18 |
| T39 $CH_3$ | H4/H7, H5/H6, $CH_3$ |
| V85 $CH_3$ | H13, H19 |
| V93 $CH_3$ | H13, H15, H19 |

**Supplementary Table 4. Contacts between hRpn13 and XL5 of the XL5-ligated hRpn13 structure measured to be within 6 Å.**

| XL5 | hRpn13 |
|---|---|
| $CH_3$ | T37 ($CH_3$); T39 (Hα, Hγ1, $CH_3$); P40 (Hγ#, Hδ#) |
| H4 | T37 ($CH_3$); V38 (Hβ, $CH_3$); T39 (HN, Hα, Hβ, Hγ1, $CH_3$); P40 (Hβ#, Hγ#, Hδ#) |
| H5 | T37 ($CH_3$); V38 (HN, Hα, Hβ, $CH_3$); T39 (HN, Hα, Hγ1, $CH_3$); P40 (Hα, Hβ#, Hγ#, Hδ#) |
| H6 | T37 ($CH_3$); V38 (HN, Hβ) |
| H7 | T37 ($CH_3$) |
| H8 | M31 (Hγ#); V38 (HN, Hα, Hβ, $CH_3$); T39 (HN, Hα); P40 (Hβ#, Hγ#, Hδ#) |
| H9 | V38 (HN, Hβ, $CH_3$); P89 (Hδ#) |
| H10 | V38 (Hβ, $CH_3$); Q87 (Hβ#); C88 (Hα, Hβ#); P89 (Hγ#, Hδ#) |
| H11 | V38 ($CH_3$); V85 ($CH_3$); Q87 (Hα, Hβ#); C88 (HN, Hα, Hβ#); P89 (Hγ#, Hδ#); S90 (HN); V93($CH_3$) |
| H12 | M31 (Hβ#, Hγ#); V38 (Hβ, $CH_3$); T39 (Hα); P40 (Hβ#, Hγ#, Hδ#); C88 (Hα, Hβ#); P89 (Hδ#); S90 (Hγ); V93($CH_3$) |
| H13 | V38 ($CH_3$); V85 (Hβ, $CH_3$); Q87 (Hβ#); C88 (HN, Hα, Hβ#); P89 (Hδ#); S90 (HN, Hβ#); V93(Hβ, $CH_3$) |
| H14 | M31 (Hγ#); V38 (Hβ, $CH_3$); 85 (Hβ, $CH_3$); Q87 (Hβ#); C88 (Hα, Hβ#); V93(Hβ, $CH_3$), F106 (Hβ#, Hδ#) |
| H15 | M31 (HN, Hα, Hβ#, Hγ#, $CH_3$); S32 (HN); L33 (Hγ); V38 ($CH_3$); V93(Hβ, $CH_3$); F106 (HN, Hα, Hβ#; Hδ#, Hε#) |
| H16 | M31 (HN, Hα, Hβ#, Hγ#); S32 (HN, Hα); L33 (HN, Hα, Hγ, $CH_3$); V38 ($CH_3$); T39 (HN); L105 (Hα); F106 (HN, Hα, Hβ#, Hδ#, Hε#, Hζ) |
| H17 | M31 (Hβ#); S32 (Hα); L33 (HN, Hα, Hβ#, Hγ, $CH_3$); V38 (Hα, $CH_3$); F106 (Hβ#, Hδ#, Hε#, Hζ) |
| H18 | L33 (Hα, Hβ#, Hγ, $CH_3$); V38 (Hα, Hβ, $CH_3$); F106 (Hδ#, Hε#, Hζ) |
| H19 | M31 (Hγ#, $CH_3$); V38 ($CH_3$); V85 (Hβ, $CH_3$); C88 (HN, Hα, Hβ#); P89 (Hδ#); S90 (HN, Hβ#, Hγ); G91 (HN); V93 (HN, Hα, Hβ, $CH_3$); F106 (Hβ#, Hδ#); W108 (Hβ#) |
| cyanide | M31 (Hγ#, $CH_3$); V38 ($CH_3$); P40 (Hβ#, Hγ#, Hδ#); C88 (Hβ#); P89 (Hδ#); S90 (HN, Hβ#, Hγ); V93 (Hβ, $CH_3$); W108 (Hβ#) |

**Supplementary Table 5. Binding affinity of XL5 derivatives for hRpn13 Pru measured by ITC.**

| 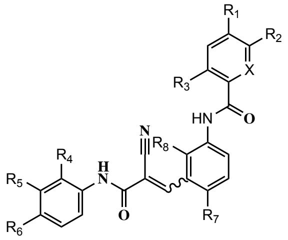 | X | R1 | R2 | R3 | R4 | R5 | R6 | R7 | R8 | NMR | ITC $K_d$ (μM) |
|---|---|---|---|---|---|---|---|---|---|---|---|
| XL5 | C | $CH_3$ | H | H | COOH | H | H | H | H | ++++++ | 1.48 ± 0.52 |
| XL23 | C | H | H | Cl | COOH | H | H | H | H | ++++ | 3.88 ± 0.43 |
| XL24 | C | -$NHCH_3$ | H | H | COOH | H | H | H | H | +++++ | 1.74 ± 0.35 |
| XL25 | C | -$NHCH_2COOH$ | H | H | COOH | H | H | H | H | +++++ | 4.12 ± 1.47 |
| XL26 | C | $CF_3$ | H | H | COOH | H | H | H | H | +++++ | 6.67 ± 1.97 |
| XL27 | N | $CH_3$ | OH | H | COOH | H | H | H | H | +++++ | 3.94 ± 1.02 |
| XL28 | C | $CH_3$ | H | H | COOH | H | -$OCH_3$ | H | H | ++++ | 3.82 ± 0.26 |
| XL29 | C | $CH_3$ | H | H | COOH | H | -$NHCH_3$ | H | H | ++++ | 7.81 ± 1.28 |
| XL30 | C | $CH_3$ | H | H | COOH | $CF_3$ | H | H | H | ++ | 12.39 ± 5.91 |
| XL31 | C | $CH_3$ | H | H | -$SO_2NH_2$ | H | H | H | H | + | NA |
| XL32 | C | $CH_3$ | H | H | COOH | H | H | $CF_3$ | H | +++ | NA |
| XL33 | C | $CH_3$ | H | H | COOH | H | H | H | -$NHCH_3$ | + | NA |

The $K_d$ value was generated by fitting ITC data to a "One Set of Sites" binding model with the Origin software. Degree of spectral changes in 2D NMR spectra is indicated by number of '+' symbols, as compared to XL5, with reduced effects symbolized by a lesser number.

**Supplementary Table 6. Six candidate sgRNAs designed by using the sgRNA Scorer 2.0 web tool.**

| ID | Target site (PAM sequence underlined) | Used for KO expts |
|---|---|---|
| 2286 | GGGCGCCTCCAACAAGTACT**TGG** | N |
| 2287 | TACTTGGTGGAGTTTCGGGC**GGG** | N |
| 2288 | GTGACTCCGGATAAGCGGAA**AGG** | Y |
| 2289 | TGACTCCGGATAAGCGGAAA**GGG** | N |
| 2290 | TCCGGATAAGCGGAAAGGGC**TGG** | Y |
| 2291 | GCTGGAAGGACAGGACGTCC**GGG** | N |

#### Supplementary Table 7. Primers for PCR.

| Primer name | Sequence |
|---|---|
| ADRM1-Amp-F | **TCCCTACACGACGCTCTTCCGATCT** CTCTCCGCGCTTTCAGGATG |
| ADRM1-Amp-R | **GTTCAGACGTGTGCTCTTCCGATCT** CAGGGACACTCACGTCTTCC |
| ADRM1-Topo-F | GCAGCCAAGACGAGAAGGTG |
| ADRM1-Topo-R | TCCGTACTCTGGGAGAGGAC |

Red = Illumina-specific primer sequence

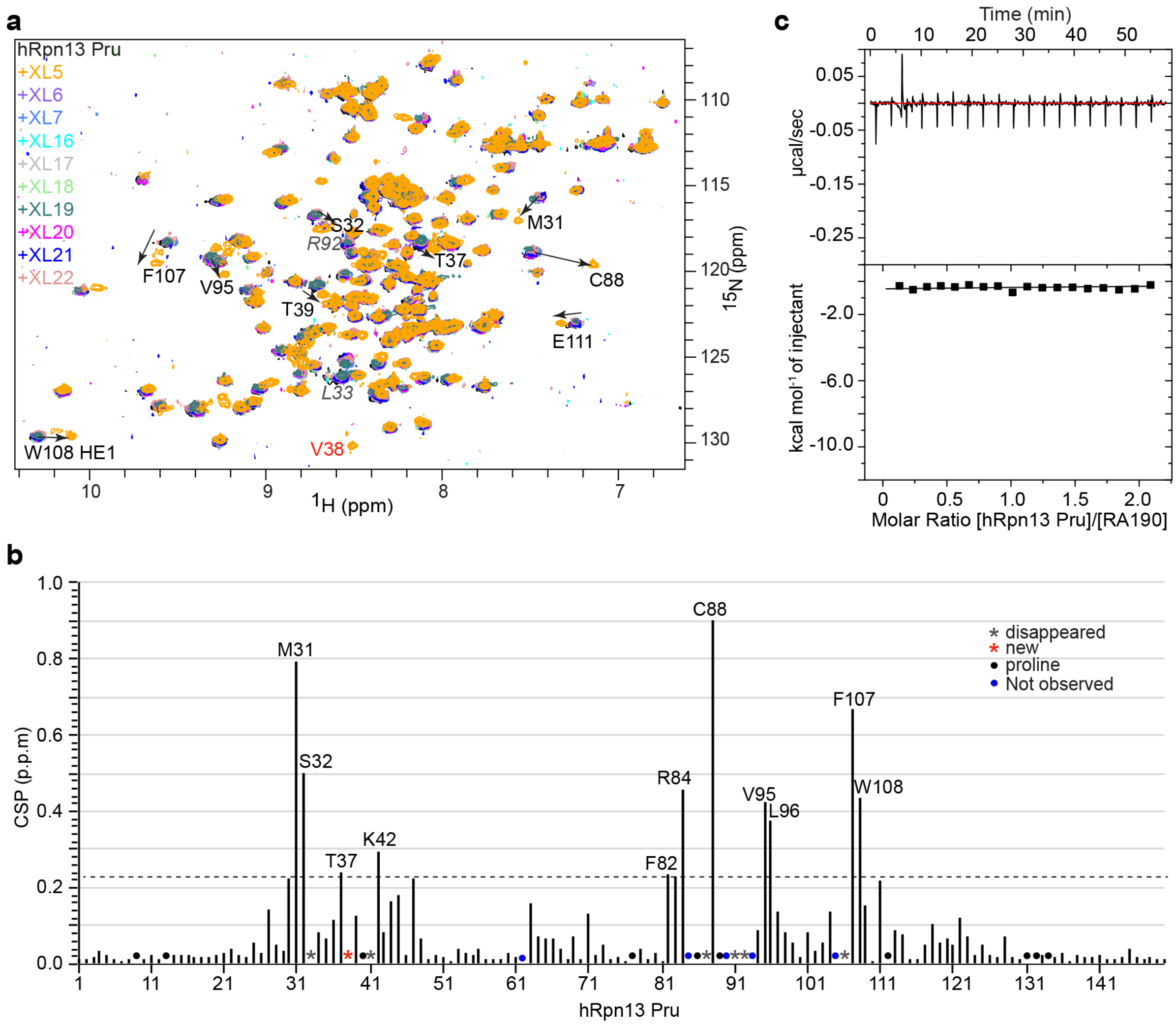


**Supplementary Fig. 1 | NMR screen identifies XL5 as binding to hRpn13. a**, $^{1}$H, $^{15}$N HSQC spectra of $^{15}$N-labeled hRpn13 Pru with addition of vehicle control DMSO (black) or 10-fold molar excess XL5 (orange), XL6 (purple), XL7 (light blue), XL16 (cyan), XL17 (grey), XL18 (green), XL19 (dark green), XL20 (magenta), XL21 (blue) or XL22 (pink). Spectra were acquired at 600 MHz and 25°C. hRpn13 signals that shift following XL5 addition are labeled and a solid arrow indicates trajectory from the free to XL5-bound state. Some signals that disappear (italicized grey) or V38 (red), which appears, following XL5 addition are also labeled. **b**, Chemical shift perturbation (CSP) values derived from the data of Fig. 1b for each hRpn13 Pru amino acid following XL5 addition. Residues with signals that disappear or appear are denoted with a grey or red star respectively. Prolines or residues not observed for free and XL5-bound hRpn13 are indicated with a black or blue dot respectively. A dashed line indicates one standard deviation above average. Residues shifted by greater than one standard deviation above average are labeled. **c**, ITC analysis of hRpn13 Pru binding to RA190. Raw ITC data (top) from titration of 200 μM hRpn13 Pru into 20 μM RA190 and binding isotherm (bottom) created by integration of the raw data.

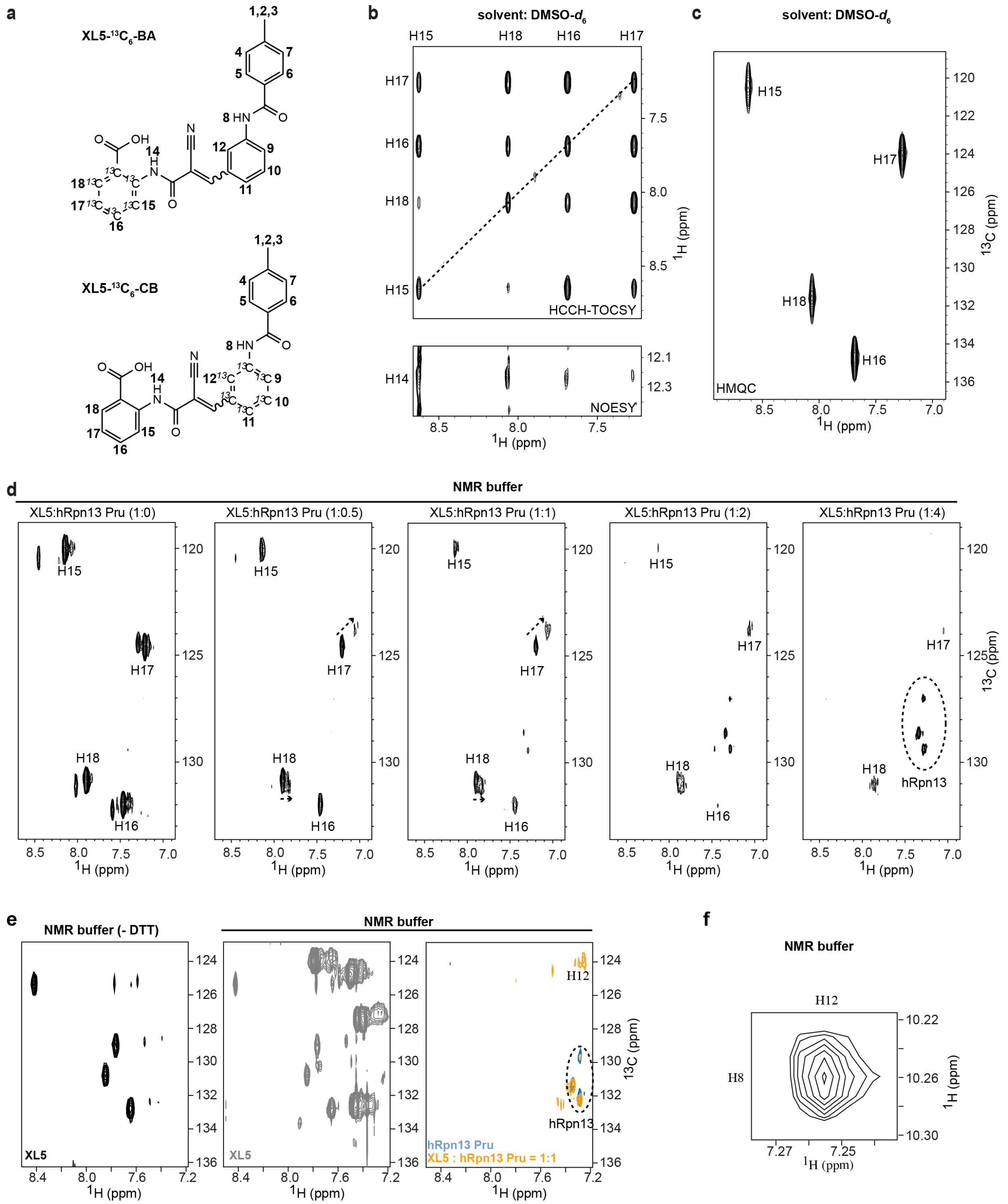


**Supplementary Fig. 2 | Chemical assignments for XL5 and after mixing with hRpn13 Pru. a**, The chemical structure of XL5 illustrating isotopic labeling of the benzoic acid carbon atoms (XL5-$^{13}C_6$-BA, top panel) or central benzene carbon atoms (XL5-$^{13}C_6$-CB, bottom panel); these labeling schemes were used for the spectra shown in **b-d** or **e-f** respectively. Hydrogen atoms are labeled with the numbers used in the text and figures. **b-c**, 2D $^{13}$C-edited HCCH-TOCSY with a 12 ms mixing time (**b**, top panel), NOESY with a 500 ms mixing time (**b**, bottom panel), and $^{1}$H, $^{13}$C HMQC spectra (**c**) recorded on 10 mM XL5-$^{13}C_6$-BA (**a**, top panel) in DMSO-$d_6$ at 25 °C. Diagonal signals in **b** are indicated by a dashed line. **d**, $^{1}$H, $^{13}$C HMQC spectra recorded on 0.1 mM XL5-$^{13}C_6$-BA (**a**, top panel) with increasing molar ratio of unlabeled hRpn13 Pru, including at 1:0 (first panel), 1:0.5 (second panel), 1:1 (third panel), 1:2 (fourth panel), and 1:4 (fifth panel) in NMR buffer. Signal shifting is indicated by a dashed arrow that extends from the free state to the hRpn13 Pru-bound state. **e**, $^{1}$H, $^{13}$C HMQC spectra recorded on 0.5 mM XL5-$^{13}C_6$-CB (**a**, bottom panel) in NMR buffer without (left panel, black) or with (middle panel, grey) DTT, or in NMR buffer and mixed with equimolar unlabeled hRpn13 Pru (right panel, orange), or on 0.5 mM unlabeled hRpn13 Pru with no XL5 (right panel, blue) in NMR buffer. **f**, Selected regions from a $^{1}$H, $^{13}$C half-filtered 2D NOESY experiment (100 ms) recorded on 0.5 mM XL5-$^{13}C_6$-CB (**a**, bottom panel) with equimolar unlabeled hRpn13 Pru in NMR buffer. Natural abundance carbon signals arising from hRpn13 Pru in **d** and **e** are indicated by a dashed oval.

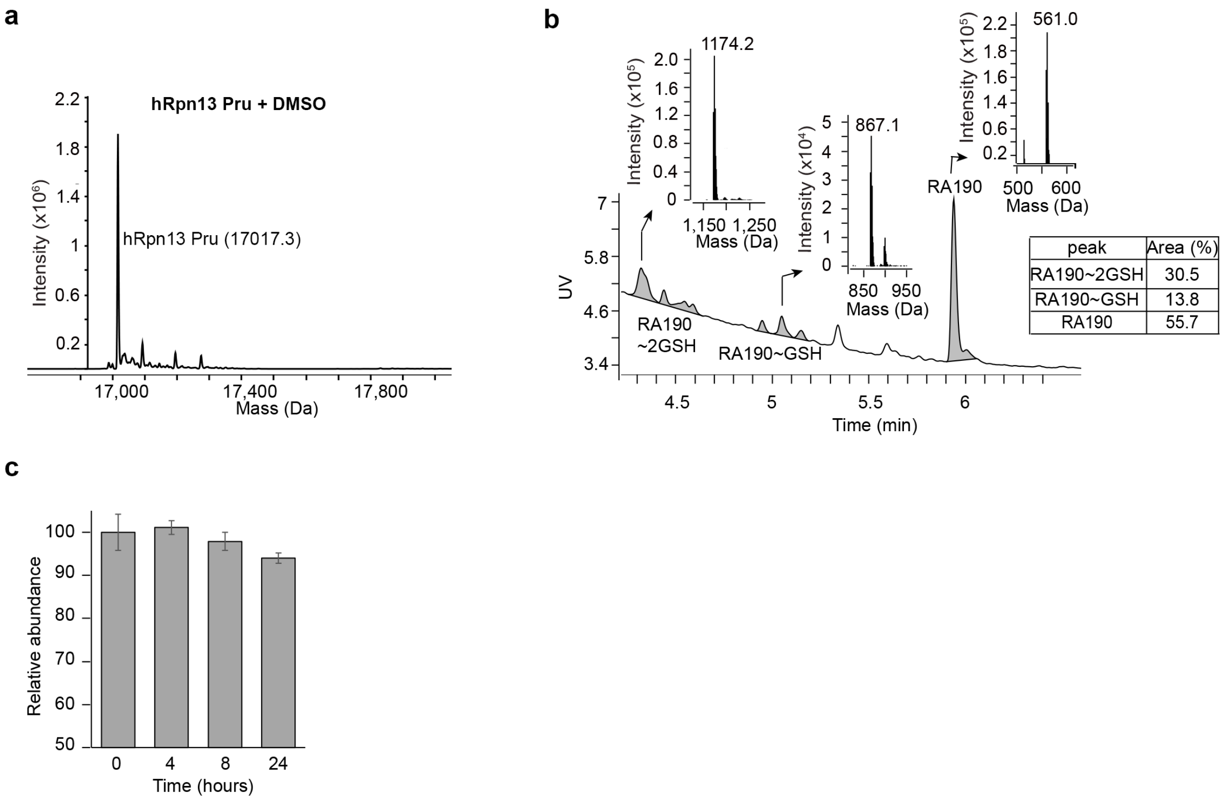


**Supplementary Fig. 3 |. LC-MS spectra examining reactivity of RA190 and XL5. a**, LC-MS analysis of 2 µM purified hRpn13 Pru (MW: 17017.3 g/mol) incubated with DMSO for 2 hours at 4°C. **b**, LC-MS analysis of 40 µM RA190 incubated with 2 mM reduced L-glutathione (GSH, MW: 307.3 g/mol) for 2 hours at 4°C. Detected GSH adducts are indicated and a table included that lists relative abundance. **c**, LC-MS analyses at indicated time points of 0.2 µM XL5 incubated at room temperature with commercially available mouse serum.

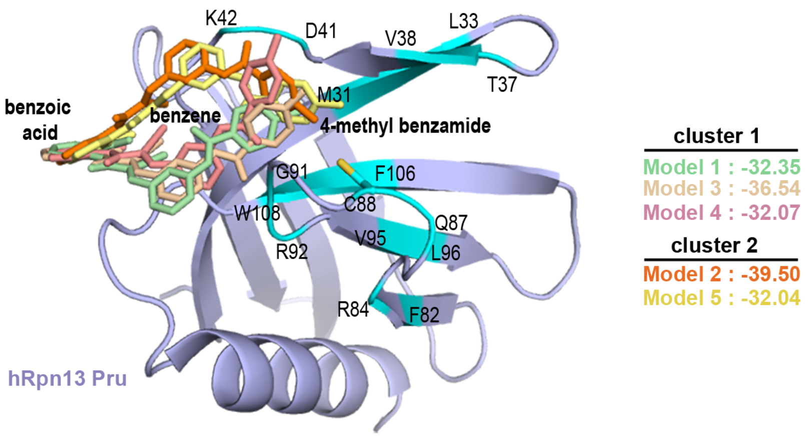


**Supplementary Fig. 4 |. Model structures of XL5 bound to hRpn13 from the *in silico* screen.** Ribbon diagram of the predicted model structures with XL5 (green in Model 1, orange in Model 2, wheat in Model 3, pink in Model 4, yellow in Model 5) bound to hRpn13 Pru (purple). Model structures are divided into two clusters based on the location of the central benzene ring, with the virtual ligand screening score calculated in ICM (Internal Coordinate Mechanics, Molsoft LLC) listed for each model. hRpn13 amino acids significantly affected by XL5 addition in Fig. 1b are highlighted in light blue as described in Fig. 1c.

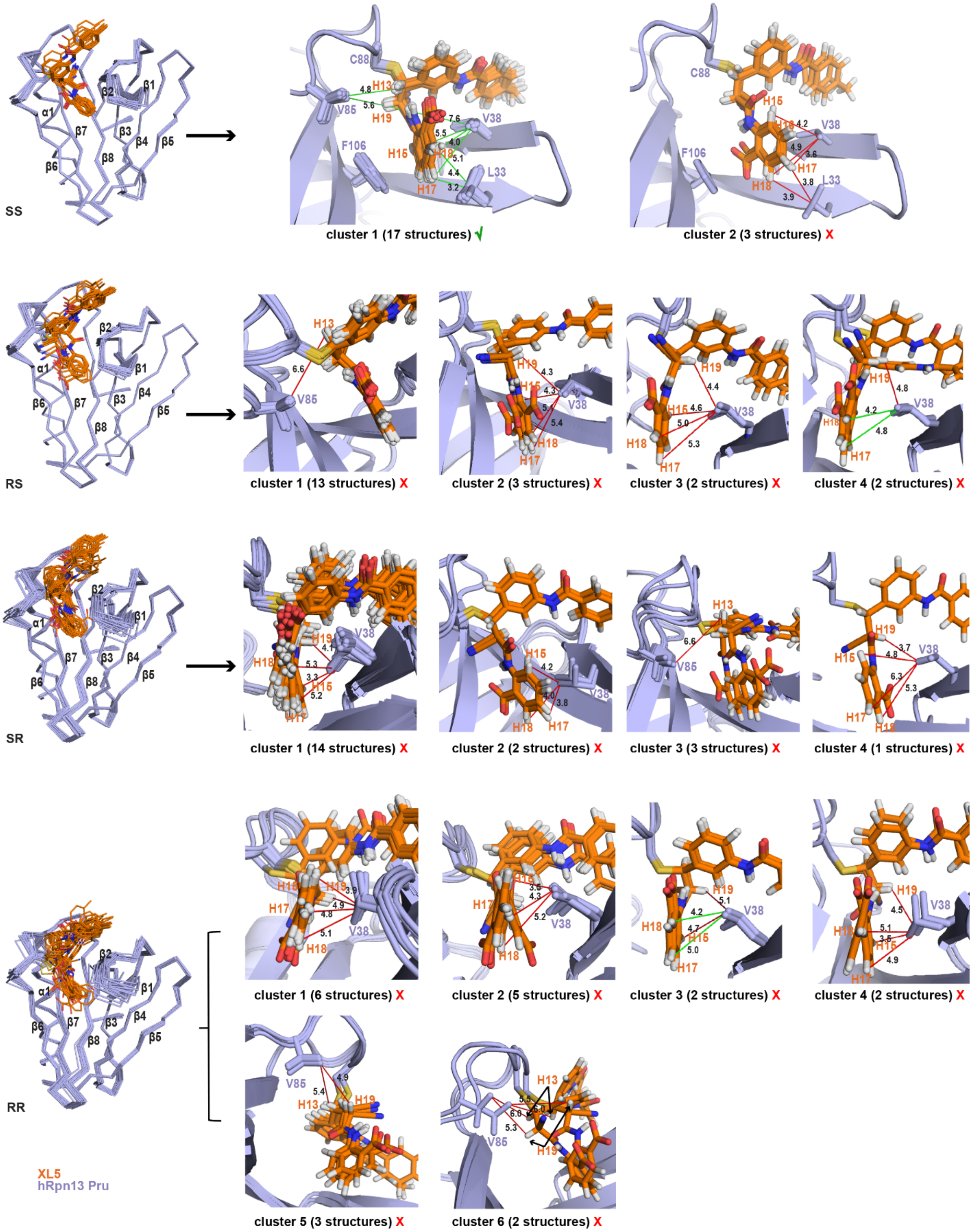


**Supplementary Fig. 5 |. Calculated structures of XL5-ligated hRpn13 Pru in different stereoisomer conformations.** hRpn13 with XL5 ligated to the C88 sulfur atom and XL5

C15 and C16 of respective SS (first panel), RS (second panel), SR (third panel) or RR (fourth panel) stereochemistry, colored as in Fig. 2g. The lowest energy structures without NOE, dihedral or torsion angle violations were clustered based on convergence. Right panels display enlarged views for each cluster centered on XL5 H13, H15, H17, H18, and H19 along with distances between atoms in Å. The number of structures within each cluster is displayed below the panel along with a green check mark (√) to indicate that the displayed interactions are supported (green lines) by the NMR data or a red x to indicate differential interactions that are not supported (red lines) by the NMR data.

**a**

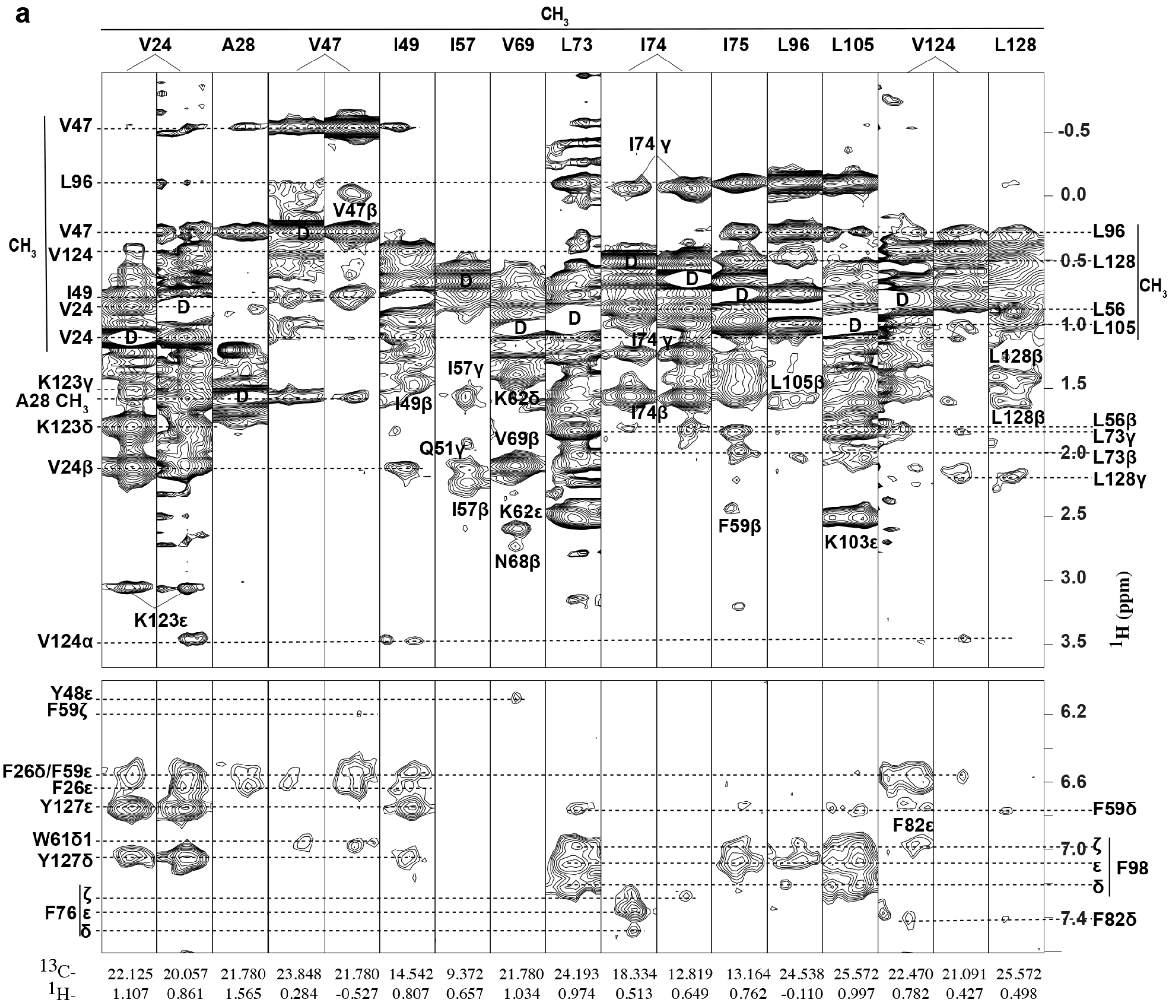


**b**

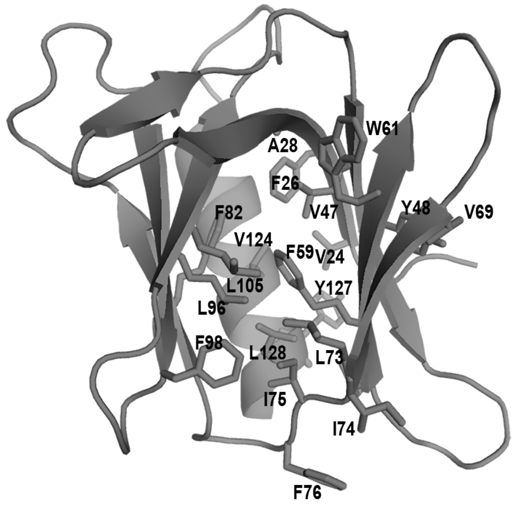


hRpn13 Pru

**Supplementary Fig. 6 | The hRpn13 Pru structure is preserved in the presence of XL5.** **a**, Selected intramolecular NOEs for XL5-bound hRpn13 Pru from a $^{1}H$, $^{13}C$ edited NOESY experiment (mixing time 100 ms) acquired with 0.4 mM $^{13}C$-labeled hRpn13 Pru and 1.2-fold molar excess unlabeled XL5. D, diagonal resonance. **b**, Residues showing NOE interactions in **a** are displayed on a ribbon diagram of free hRpn13 Pru (grey, PDB 5IRS). The amino acids displayed in **a** belong to the structural core and show the expected interactions for structural integrity.

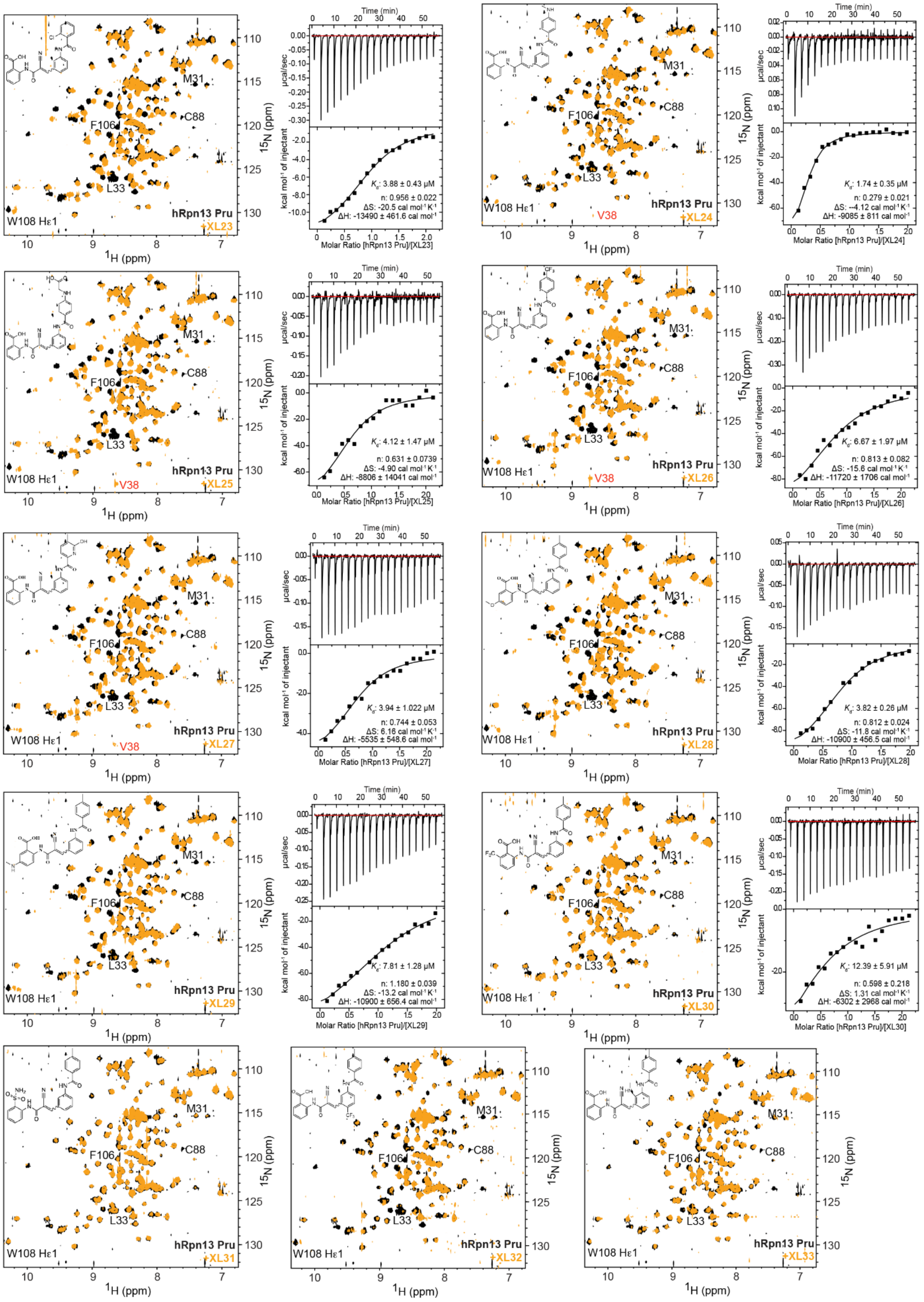
hRpn13 Pru
XL23
M31
C88
F106
L33
W108 Hε1
$^{1}$H (ppm)
$^{15}$N (ppm)
Time (min)
μcal/sec
kcal mol$^{-1}$ of injectant
$K_d$: 3.88 ± 0.43 μM
n: 0.956 ± 0.022
ΔS: -20.5 cal mol$^{-1}$ K$^{-1}$
ΔH: -13490 ± 461.6 cal mol$^{-1}$
Molar Ratio [hRpn13 Pru]/[XL23]
XL24
V38
$K_d$: 1.74 ± 0.35 μM
n: 0.279 ± 0.021
ΔS: --4.12 cal mol$^{-1}$ K$^{-1}$
ΔH: -9085 ± 811 cal mol$^{-1}$
Molar Ratio [hRpn13 Pru]/[XL24]
XL25
$K_d$: 4.12 ± 1.47 μM
n: 0.631 ± 0.0739
ΔS: -4.90 cal mol$^{-1}$ K$^{-1}$
ΔH: -8806 ± 14041 cal mol$^{-1}$
Molar Ratio [hRpn13 Pru]/[XL25]
XL26
$K_d$: 6.67 ± 1.97 μM
n: 0.813 ± 0.082
ΔS: -15.6 cal mol$^{-1}$ K$^{-1}$
ΔH: -11720 ± 1706 cal mol$^{-1}$
Molar Ratio [hRpn13 Pru]/[XL26]
XL27
$K_d$: 3.94 ± 1.022 μM
n: 0.744 ± 0.053
ΔS: 6.16 cal mol$^{-1}$ K$^{-1}$
ΔH: -5535 ± 548.6 cal mol$^{-1}$
Molar Ratio [hRpn13 Pru]/[XL27]
XL28
$K_d$: 3.82 ± 0.26 μM
n: 0.812 ± 0.024
ΔS: -11.8 cal mol$^{-1}$ K$^{-1}$
ΔH: -10900 ± 456.5 cal mol$^{-1}$
Molar Ratio [hRpn13 Pru]/[XL28]
XL29
$K_d$: 7.81 ± 1.28 μM
n: 1.180 ± 0.039
ΔS: -13.2 cal mol$^{-1}$ K$^{-1}$
ΔH: -10900 ± 656.4 cal mol$^{-1}$
Molar Ratio [hRpn13 Pru]/[XL29]
XL30
$K_d$: 12.39 ± 5.91 μM
n: 0.598 ± 0.218
ΔS: 1.31 cal mol$^{-1}$ K$^{-1}$
ΔH: -6302 ± 2968 cal mol$^{-1}$
Molar Ratio [hRpn13 Pru]/[XL30]
XL31
XL32
XL33

**Supplementary Fig. 7 |. NMR and ITC analyses of hRpn13 binding to XL5 derivatives.** $^{1}H$, $^{15}N$ HSQC spectra of 20 μM $^{15}N$-labeled hRpn13 Pru with addition of vehicle control DMSO (black) or 10-fold molar excess XL5 derivative XL23-XL33 (orange), as indicated, and accompanying ITC data for XL5 derivatives XL23-XL30. NMR spectra were acquired at 10°C and 600 MHz for XL23-XL32 and 800 MHz for XL33. The signal for hRpn13 V38 is labeled in red when it appears. XL5 binding residues hRpn13 M31, L33, C88, F106, W108 are labeled. ITC analyses are presented to the right of the corresponding NMR data when available with the top panel plotting raw ITC data and the bottom panel displaying the binding isotherm generated by integrating the raw data. ITC titrations were performed with 200 μM hRpn13 Pru injected into 20 μM XL5 derivative. The data was fit to a “One Set of Sites” binding model with the indicated thermodynamic values by using the Origin software. The chemical structure of each XL5 derivative is included within the NMR spectra.

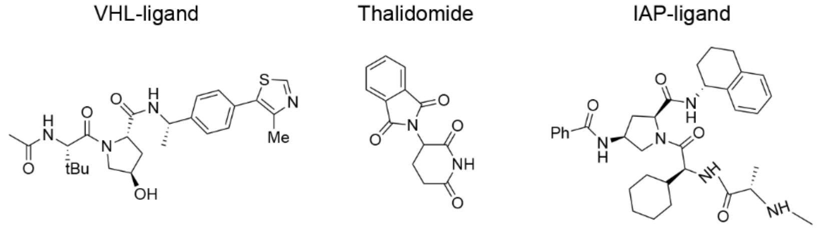


**Supplementary Fig. 8 |. Chemical structures of PROTACs fused with XL5, including VHL-ligand, thalidomide and IAP-ligand.**

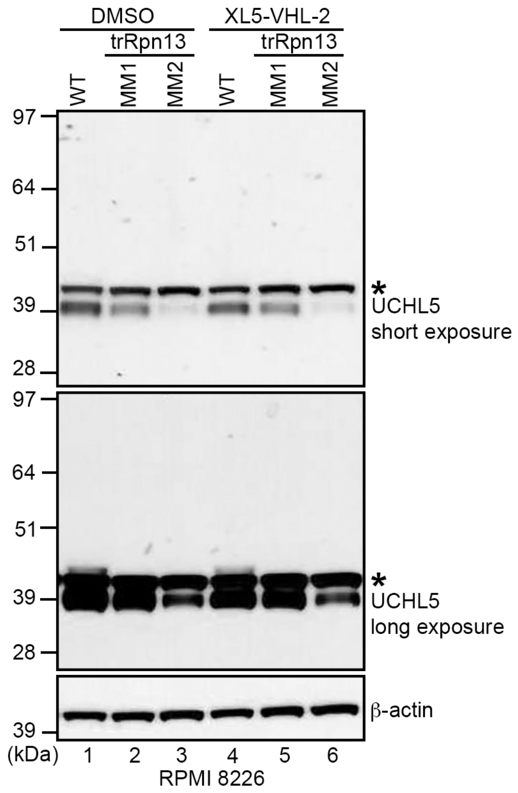


**Supplementary Fig. 9 |. UCHL5 does not appear to be targeted by XL5-VHL-2.** Immunoblots of whole cell lysate from RPMI 8226 WT, trRpn13-MM1, or trRpn13-MM2 cells treated for 24 hours with 40 μM XL5-VHL-2 with comparison to DMSO (vehicle control) immunoprobing for UCHL5 (short and long exposure), or β-actin (as a loading control, bottom panel). A black asterisk indicates an unknown band that may be non-specific.

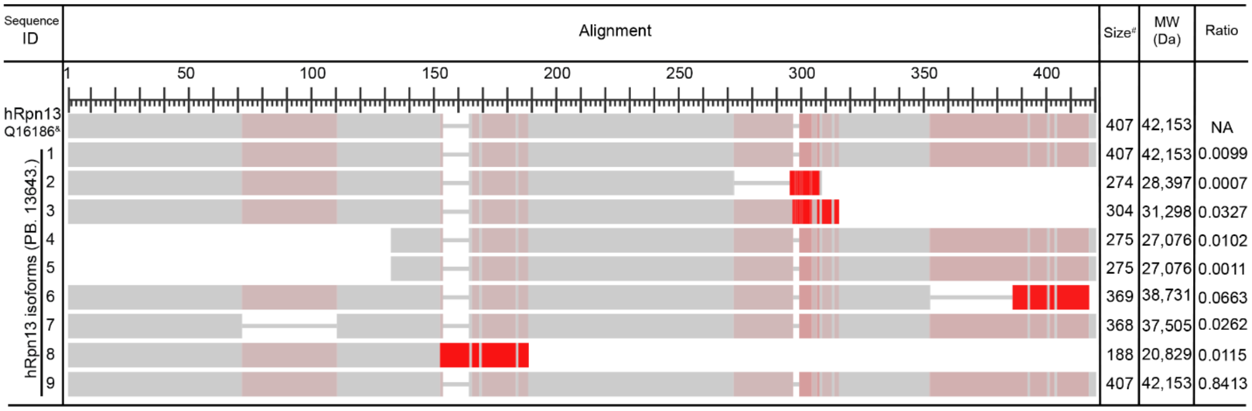


### Size provided as amino acid number

& UniProtKB number of hRpn13

**Supplementary Fig. 10 |. Protein sequence alignment of hRpn13 isoforms in RPMI 8226 WT cells.** The protein sequence of hRpn13 isoforms translated from mRNA transcripts identified by mRNA PacBio were aligned with Protein Blast (https://blast.ncbi.nlm.nih.gov/BlastAlign.cgi) and colored based on frequency-based amino acid variance. Darker shades of red indicate greater difference from residues in other rows of the alignment at that position. Included is sequence ID, size (number of amino acids), molecular weight (MW) and ratio of hRpn13 mRNA expression for each isoform as quantified by Illumina. The mRNA samples were isolated from RPMI 8226 WT cells and analyzed in triplicate.
